# Bacteriophage P22 virus-like particles as nanoscale protein scaffolds for plant synthetic biology

**DOI:** 10.64898/2026.05.14.725278

**Authors:** Maxim D. Harding, Mark A. Jackson, Edward K. Gilding, David J. Craik, Frank Sainsbury, Nicole Lawrence

## Abstract

Advancing the utility of plant synthetic biology requires the continued development of protein engineering tools. Self-assembling protein compartments, such as virus-like particles (VLPs), provide versatile scaffolds for synthetic biology. However, few plant-expressed VLPs have demonstrated broad amenability to protein engineering, restricting their applications to specific contexts. Here, the *Salmonella typhimurium* bacteriophage P22 VLP is explored as a novel protein scaffold for plant synthetic biology, demonstrating its application in a eukaryote for the first time. Through transient expression in the biofactory plant *Nicotiana benthamiana*, the capacity for P22 VLPs to correctly assemble and selectively encapsulate recombinant protein cargo is demonstrated. The durability of this protein scaffold is explored, through co-encapsulation of multiple cargo protein species and by encapsulation through direct fusion to the P22 coat protein. Finally, the ability to simultaneously program cargo encapsulation and external protein display on P22 VLPs *in vivo* is demonstrated through SpyTag/SpyCatcher-mediated protein conjugation. This work demonstrates the broad utility of P22 VLPs as nanoscale protein scaffolds for plant synthetic biology.

## Introduction

Synthetic biology relies on the ability to engineer cells at the molecular level. Protein scaffolds comprised of self-organising units that assemble into functional complexes are valuable tools for synthetic biology, capable of selectively recruiting proteins to distinct locations within a cell. For example, when reconstituting or engineering novel metabolic pathways, protein scaffolds can be used to co-localise enzymes to achieve enhanced metabolic flux through proximity (Dueber et al., 2009; Sweetlove and Fernie, 2018). Traditionally, naturally evolved protein scaffolds have been engineered by replacing the native tethered enzyme with heterologous enzymes of choice (Agapakis et al., 2012). These scaffolds can include soluble protein interaction domains (Dueber et al., 2009), or membrane proteins tethered to organelles, the cytoplasm, or lipid droplets (Behrendorff et al., 2020). Many of these scaffolding approaches have been employed *in planta* for optimising recombinant product biosynthesis (Harding and Sainsbury, 2025). More recently, a range of bioengineered protein scaffolds have been developed, exploiting unique functionalities such as phase-separation within biomolecular condensates (Hilditch et al., 2024; Lindström Battle et al., 2025; Qian et al., 2022), and encapsulation within or display outside of self-assembling protein compartments (Cheah et al., 2023; Giessen and Silver, 2016; Hartzell et al., 2020; Lau et al., 2018; Su et al., 2023).

Virus-like particles (VLPs) are a class of self-assembling protein scaffold, rich with structural and functional diversity that is advantageous for applications in metabolic engineering and therapeutic development (McNeale et al., 2023a; Su et al., 2023). Bacteriophage P22 VLPs have been employed as synthetic scaffolds due to their stability, large internal volume, and corresponding encapsulation capacity (Esquirol et al., 2022; McNeale et al., 2023a; O’Neil et al., 2011). Comprised of 420 coat proteins (CPs) and approximately 100-330 scaffold proteins (SPs), P22 VLPs assemble *in vivo* to form 58 nm diameter T=7 icosahedral capsids (Thuman-Commike et al., 1998). During particle assembly, the SPs accumulate within the interior of P22 VLPs via ionic interactions with the inward facing surface of self-assembling CPs. Through genetic fusion to either the N- or C-terminus of the SP, cargo such as enzymes, peptides, and other proteins of interest can be encapsulated within P22 VLPs (Esquirol et al., 2022; Harding et al., 2025; Patterson et al., 2012). Encapsulation of multiple cargo species can be achieved by tandem fusion to a single SP (Patterson et al., 2014), or by co-expression of different cargo-SP fusions (McNeale et al., 2023b), enabling co-localisation of subsequent enzymes from metabolic cascades. P22 VLPs are also amendable to external scaffolding applications due to the C-terminus of the CP being externalised on assembled VLPs. For example, genetic fusion of respective sequences to the CP C-terminus has allowed external functionalisation of P22 VLPs via SpyTag/SpyCatcher mediated isopeptide bond formation (Kim et al., 2019), and enzyme-mediated ligation using sortase A (Patterson et al., 2017). The flexibility of P22 VLPs for both encapsulation and external display make them excellent scaffolds for protein engineering applications. However, despite the potential of self-assembling protein scaffolds like VLPs for engineering applications in eukaryotes (Lau et al., 2018), to date P22 VLPs have only been expressed in *Escherichia coli*.

The biofactory plant *Nicotiana benthamiana* has proven to be highly capable of producing a range of different VLPs at commercial scale. For example, the SARS-CoV-2 spike protein and influenza haemagglutinin, which effectively form lipid-enveloped VLPs (D’Aoust et al., 2008; Jung et al., 2022) when targeted to the apoplast, have both reached clinical application (Ward et al., 2021; Ward et al., 2020). Although only lipid-enveloped VLPs expressed in plants have reached clinical settings as vaccines to-date, non-enveloped VLPs are also showing promise and may have the advantage of broader applicability as protein scaffolds. Examples of non-enveloped VLPs with therapeutic potential include cowpea mosaic virus VLPs, which have been shown to induce systemic antitumour immunity upon injection in mouse models of metastatic melanoma (Lizotte et al., 2016), and expression of adeno-associated viral particles in *N. benthamiana* for use in gene therapy applications, which was recently patented by the US-based startup Cirsium Bioscience (Gibbs and Connors, 2021).

Non-enveloped VLP protein scaffolds can be categorised according to their functionality, including as scaffolds engineered for programmable cargo encapsulation, or as scaffolds engineered for external protein display. For example, bluetongue virus VLPs transiently expressed in the cytosol of *N. benthamiana* have been engineered to encapsulate herpes simplex virus thymidine kinase as a pro-drug converting enzyme for cancer therapy (Brillault et al., 2017; Thuenemann et al., 2021). Recently, one study reported the expression of MS2 bacteriophage CP in the tobacco chloroplast for applications in RNA interference against cotton bollworm (Jiang et al., 2024), indicating the ability to also use VLP components for the protection of nucleic acids *in planta*. Engineering VLP scaffolds for external protein display has also been highly effective in *N. benthamiana*. Hepatitis B core (HBc) VLPs have been used as scaffolds for external protein display through SpyTag/SpyCatcher-mediated isopeptide bond formation, in both the cytosol and endoplasmic reticulum, via engineering of SpyCatcher within an exposed loop of HBc VLPs and co-expression of either GFP or the human immunodeficiency virus capsid protein P24 fused to SpyTag (Peyret et al., 2020). Additionally, plant expressed HBc VLPs, tomato bushy stunt virus, and potato virus X have been engineered as scaffolds for external functionalisation via E-coil and K-coil mediated dimerization (Sator et al., 2024). Some VLPs, such as grapevine fanleaf virus VLP previously expressed in *N. benthamiana* (Belval et al., 2016), are amendable to engineering both encapsulation and external protein display. VLPs with such flexibility as protein scaffolds can be applied to diverse engineering contexts and are thus valuable synthetic biology tools.

Here, we explore the utility of P22 VLPs as broadly applicable protein engineering scaffolds in a eukaryote for the first time, capable of both programmable encapsulation and external protein display in *N. benthamiana*. First, we demonstrate the capacity for *N. benthamiana* to produce correctly assembled P22 VLPs with up to two co-encapsulated cargo proteins exploiting non-covalent interactions with the SP. Next, we explore the ability to encapsulate toxic peptide cargo, unable to be packaged as SP-fusions, using direct fusion to the CP N-terminus. Finally, we employ the SpyTag/SpyCatcher system to functionalise the exterior of P22 VLPs with proteins of interest *in vivo*.

## Results and Discussion

### Bacteriophage P22 VLPs assemble and package selective protein cargo *in planta*

To first assess whether P22 VLPs can assemble *in planta*, the P22 CP and the fluorescent reporter protein mRUBY3 fused to the C-terminus of the P22 SP (SP-mRUBY3) were incorporated into independent pEAQ-HT transient expression vectors (Figure 1A, Table S1) (Sainsbury et al., 2009). These constructs were assembled without a signal peptide such that recombinant proteins should accumulate in the plant cytosol. Given the demonstrated success of co-infiltration of multiple *Agrobacterium tumefaciens* strains harbouring independent expression vectors (Alamos et al., 2025; Carlson et al., 2023), it was reasoned that co-infiltrating a 1:1 ratio of *A. tumefaciens* harbouring pEAQ P22 CP and pEAQ SP-mRUBY3 respectively would facilitate co-expression of these proteins in the same cell. To assess protein expression, VLP assembly, and SP-mRUBY3 cargo packaging, protein extracts from transient expressions (n=5) were sedimented by ultracentrifugation at 150,000 g and analysed by sodium dodecyl-sulfate polyacrylamide gel electrophoresis (SDS-PAGE) (Figure 1B). Coomassie staining of the gel showed the presence of a strong protein band below 50 kDa and a less prominent band below 40 kDa. In-gel tryptic digest followed by liquid chromatography tandem mass spectrometry (LC-MS/MS) analysis confirmed the identity of both species as being the P22 CP and SP-mRUBY3 respectively (Table S2), demonstrating successful transient expression.

**Figure 1.**
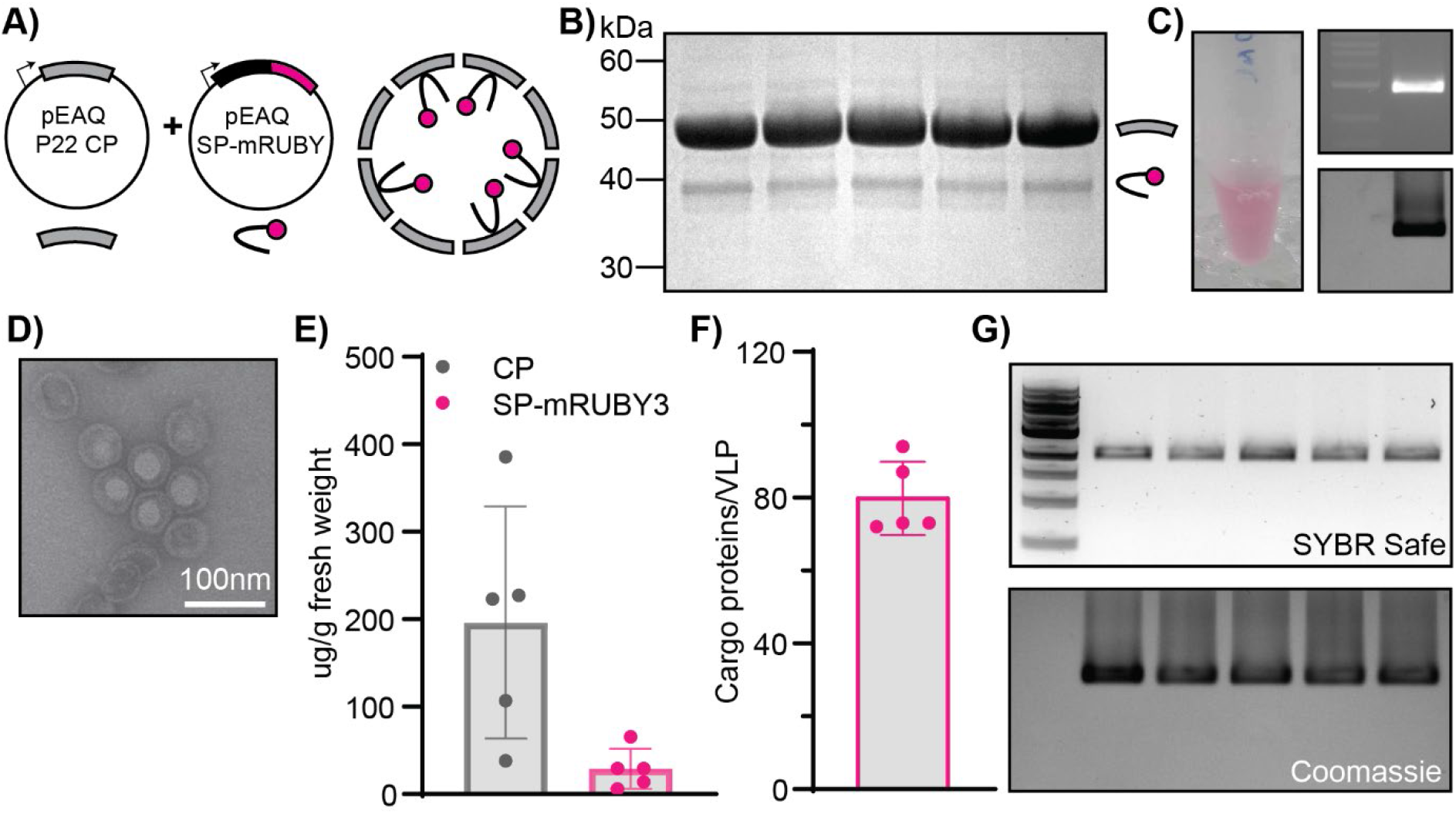
P22 VLP assembly and single cargo packaging. **A)** Cartoon illustration of P22 CP and SP-mRUBY3 expression vectors and packaged P22 VLP. **B)** Coomassie stained SDS-PAGE gel for purified P22 SP-mRUBY3 VLPs (n=5). Identity of protein band species are depicted on the right. **C)** Visual pink fluorescence from ultracentrifuged VLP extract and corresponding native agarose gel. Images of protein sample visualised with fluorescence (top) and Coomassie stain (bottom) on a UV gel documentation system. **D)** Negative stain TEM of purified P22 SP-mRUBY3 VLPs. Scale bar is 100 nm. **E)** Calculated yields of P22 CP and SP-mRUBY3 in µg/g plant fresh weight. **F)** Cargo loading stoichiometries for SP-mRUBY3 per VLP (n=5). **G)** Native agarose gel showing nucleic acid packaging by SYBR Safe (nucleic acid) stain and consequent Coomassie stain. Lane 1 is a 1 kb nucleic acid ladder.

The proteins were harvested by ultracentrifugation to separate large protein complexes such as ∼19 MDa P22 VLPs from smaller, soluble proteins. The pink fluorescence of the harvested protein pellet (Figure 1C) suggests that SP-mRUBY3 was successfully packaged within P22 VLPs. Native agarose gel electrophoresis (NAGE) of the purified sample further supports SP-mRUBY3 packaging, as the visualised fluorescent signal corresponded to the Coomassie stained protein species (Figure 1C). Furthermore, the presence of a single band in the native agarose gel indicates that the P22 CP and SP-mRUBY3 proteins are complexed as a single migrating species, and negative stain transmission electron microscopy (TEM) confirmed correctly assembly of P22 VLPs packaged with SP-mRUBY3 cargo (Figure 1D).

Average recombinant protein yields were calculated to be 196 µg/g and 28 µg/g fresh weight for the P22 CP and SP-mRUBY3 respectively (Figure 1E). The molar ratio reflects cargo packaging efficiency, as only packaged SP-mRUBY3 sediments with the VLPs. The stoichiometry of packaged SP-cargo to CP has been demonstrated to range from 45 to 275 cargo proteins per VLP in *E. coli* (Esquirol et al., 2022). Average loading of SP-mRUBY3 in plant expressed P22 VLPs was 80 cargo proteins/VLP – assuming correct P22 particle morphology comprising 420 CP/VLP (Figure 1F). Interestingly, SYBR Safe staining of the purified protein samples following NAGE revealed packaging of nucleic acid within the cytosolically expressed P22 VLPs, with the visualised nucleic acid band corresponding to the Coomassie stained P22 VLP protein band (Figure 1G). This observation suggests that, like other VLPs recombinantly expressed in plants, P22 VLPs produced in *N. benthamiana* non-specifically package nucleic acids in addition to the recombinant protein cargo.

To validate that the P22 VLP was produced by *N. benthamiana*, and not the *A. tumefaciens* used in the transformation process (Gelvin, 2003), intron 2 from potato ST-LS1 (referred to herein as ST-LS1) was inserted into the P22 CP sequence (CP ST-LS1). This intron has been demonstrated to increase the stability of a pepper mottle virus infectious clone in *E. coli*, while being correctly spliced out in *N. benthamiana* to enable viral infection (Tran et al., 2019). Insertion of the ST-LS1 intron at 198 bp within the P22 CP coding sequence was designed to terminate translation in *A. tumefaciens* with an opal stop codon, located 8 codons after lysine 66 of the P22 CP (Figure S1A, Table S3). With this design, only correctly spliced CP transcripts would facilitate translation of full-length CP and consequent assembly of P22 VLPs – a process that could not occur in *A. tumefaciens* due to the absence of splicing machinery. Co-expression of the modified CP ST-LS1 construct with SP-mRUBY3 in *N. benthamiana* yielded full-length P22 CP, and correctly assembled P22 VLPs with SP-mRUBY3 packaged (Figure S1B, Table S2), confirming the production of P22 VLPs by *N. benthamiana*. Interestingly, whereas the average yield of the purified CP from the CP ST-LS1 expressions was similar to those with the wild-type CP, the yield of SP-mRUBY3 was 3-fold greater (86 µg/g fresh weight) with 188 cargo proteins loaded per VLP (Figure S1C, D). It is possible that the additional kinetic step of ST-LS1 intron splicing decreased the rate of P22 CP expression and particle assembly, thereby enabling more SP-mRUBY3 cargo to accumulate for packaging. However, further quantitative work would be required to confirm this hypothesis, which may reveal a useful construct design consideration for multicomponent recombinant products in plants where control over the rate of expression is useful.

### Co-encapsulation of two fluorescent protein cargoes in P22 VLPs

P22 VLPs have been employed in synthetic biology as enzyme nanoreactors due to their capacity to co-encapsulate multiple cargo species, such as sequential enzymes from catalytic cascades (Jordan et al., 2016; McNeale et al., 2023b; Patterson et al., 2014; Wang et al., 2023). To determine whether two distinct cargoes can be co-encapsulated within the P22 VLPs *in planta*, the *A. tumefaciens* strains harbouring the wild-type P22 CP and SP-mRUBY3 were co-infiltrated together with a third strain harbouring the fluorescent protein mCLOVER3 fused to the C-terminus of P22 SP (SP-mCLOVER3), at a 1:1:1 ratio (Figure 2A). Fluorescent imaging of a single *N. benthamiana* leaf 5-days after infiltration with each fluorescent protein *A. tumefaciens* strain along with the P22 CP strain confirmed successful expression and fluorescence of both SP-mRUBY3 and SP-mCLOVER3 (Figure 2B, Figure S2). The spot infiltrations for P22 SP-mCLOVER3 and P22 SP-mRUBY3 (Figure 2B (ii) and (iii), respectively) demonstrate successful expression of the individual fluorescent proteins, and the co-infiltrated spot (Figure 2B (iv)) showed co-localised expression of the fluorescent proteins.

**Figure 2.**
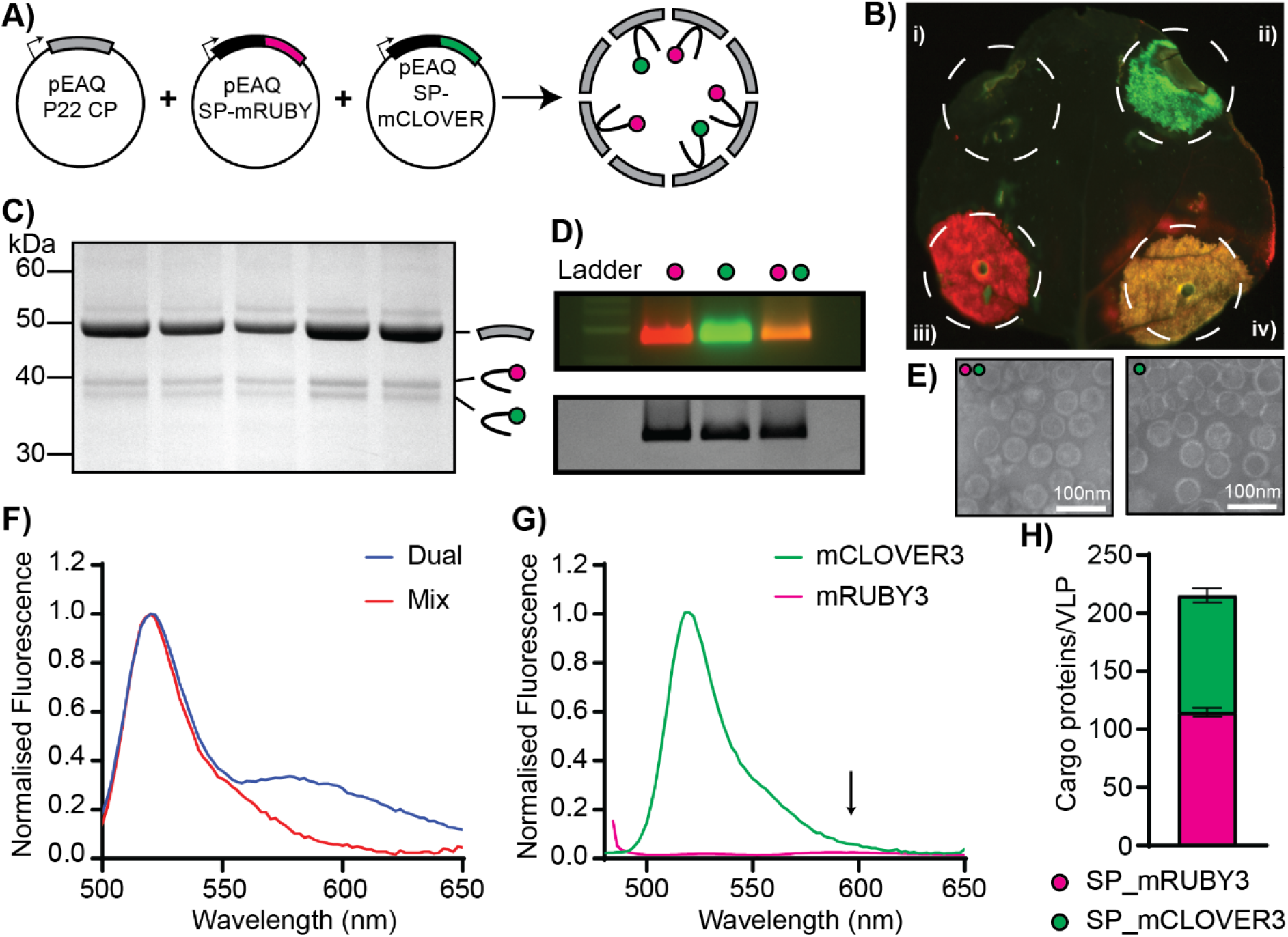
Dual loading of P22 VLPs with two fluorescent protein cargoes. **A)** Cartoon illustration of pEAQ expression vectors harbouring P22 CP, SP-mRUBY3, and SP-mCLOVER3, and a schematic of dual loaded P22 VLP harbouring both fluorescent protein cargoes. **B)** Fluorescent image of an *N. benthamiana* leaf 5-days post infiltration with (i) pEAQ P22 CP, (ii) pEAQ P22 CP + pEAQ SP-mCLOVER3, (iii) pEAQ P22 CP + pEAQ SP-mRUBY3, and (iv) pEAQ P22 CP + pEAQ SP-mCLOVER3 + pEAQ SP-mRUBY3. **C)** Coomassie stained SDS-PAGE of ultracentrifuged purified P22 VLPs with dual packaged SP-mRUBY3 and SP-mCLOVER3 (n=5). **D)** Native agarose gel (above) imaged with fluorescence channels for mCherry (red) and GFP (green), and (below) imaged after overnight staining with Coomassie. Lane 1 contains 1kb DNA ladder, lane 2 (pink dot) contains P22 SP-mRUBY3 VLPs, lane 3 (green dot) contains P22 SP-mCLOVER3 VLPs, and lane 4 (green and pink dots) contains dual loaded P22 SP-mRUBY3 + SP-mCLOVER3 VLPs. **E)** Negative stain TEM of dual loaded P22 VLPs and P22 SP-mCLOVER3 VLPs. **F)** Fluorescent spectra (λ_ex_ = 450 nm) of dual loaded P22 VLPs vs a mix of P22 SP-mRUBY3 and P22 SP-mCLOVER3 VLPs, normalised at mCLOVER3 emission (518 nm). **G)** Bleed-through from mCLOVER3 emission and direct excitation of mRUBY3 by excitation at 450 nm. Arrow indicates emission maxima for mRUBY3. **H)** Cargo loading stoichiometries for dual loaded P22 (n=5).

Analysis of the Coomassie stained SDS-PAGE and tryptic digest LC-MS/MS confirmed the presence of both SP-mRUBY3 and SP-mCLOVER3 cargo in the ultracentrifuged protein extracts from the co-infiltrated expressions (n=5) (Figure 2C, Table S2). Furthermore, fluorescent imaging (top) and Coomassie staining (bottom) of the native agarose gel confirmed the presence of both fluorescence protein cargoes in the co-infiltrated VLP extracts (Figure 2D). Correct assembly and morphology of P22 VLPs with SP-mCLOVER3 alone, or together with SP-mRUBY3, was confirmed with negative stain TEM (Figure 2E) and nano flow cytometry (Figure S3).

To confirm that SP-mRUBY3 and SP-mCLOVER3 were co-encapsulated within P22 VLPs (herein referred to as dual-loaded VLPs), a Förster resonance energy transfer (FRET) experiment was conducted. mCLOVER3 acts as a FRET donor for mRUBY3 (Bajar et al., 2016), when the fluorescent pairs are within a 1-10 nm distance of one another (Shrestha et al., 2015). Therefore, it was hypothesised that only dual-loaded P22 VLPs would facilitate FRET upon mCLOVER3 excitation (leading to mRUBY3 excitation through proximity), whereas a mixture of P22 VLPs packaged with either SP-mRUBY3 or SP-mCLOVER3 independently would not excite mRUBY3. Indeed, upon excitation of mCLOVER3 at 450 nm, only the dual-loaded P22 VLPs emitted a fluorescence intensity signal that corresponded to mRUBY3 emission (590 nm), and the mixed sample only depicted a peak corresponding to mCLOVER3 emission (518 nm) (Figure 2F). Moreover, direct excitation (450 nm) of P22 VLPs independently packaged with SP-mCLOVER3 or SP-mRUBY3, produced a strong fluorescence signal for mCLOVER3, but negligible spectral bleed-through for mRUBY3 (Figure 2G). This result confirms the dual loading of both SP-mRUBY3 and SP-mCLOVER3 cargo in P22 VLPs when co-expressed in *N. benthamiana*. Packaging of the fluorescent proteins was determined by SDS-PAGE densitometric analysis to be 114 and 100 molecules of SP-mRUBY3 and SP-mCLOVER3 per P22 VLP on average (Figure 2H). The ability to program the co-encapsulation of two different protein cargoes within P22 VLPs provides a new tool for subcellular organisation which could be applied for metabolic engineering in *N. benthamiana*.

### Expanding *N. benthamiana’s* biosynthetic range by encapsulation of toxic cargo

Engineering *N. benthamiana* to produce high-value compounds like industrial precursors of chemotherapy drugs paclitaxel and etoposide (Lau and Sattely, 2015; McClune et al., 2025), peptide therapeutics such as exenatide (Akter et al., 2022; Choi et al., 2014; Kwon et al., 2013), and a wide range of protein biologics (Eidenberger et al., 2023) have been thoroughly investigated over the past 20 years. However, limitations remain for the heterologous production of products with undesirable toxicity to the plant host. One notable example of this is the production of highly charged peptide therapeutics, such as antimicrobial peptides (AMPs), which often cause a necrotic phenotype in host plant tissues resulting in an inability to purify recombinant product (Ghidey et al., 2020). Recent work addressing this challenge has successfully demonstrated that for some AMPs genetic fusion to the small ubiquitin-like modifier (SUMO) protein can alleviate toxicity to the plant host when expressed stably in the plastid (Hoelscher et al., 2022) or transiently in the cytosol (Chaudhary et al., 2023). It was hypothesised that encapsulation of charged bioactive peptides might also alleviate toxicity to the plant host by sequestering them in the internal lumen of P22 VLPs (Harding et al., 2025). The antiplasmodial peptide PDIP (Platelet factor 4-derived internalisation peptide) (Lawrence et al., 2018; Lawrence et al., 2024) was selected as a representative charged bioactive peptide to explore whether encapsulation within VLPs could facilitate accumulation in *N. benthamiana* as previously demonstrated in *E. coli* (Harding et al., 2025).

As described for the individual fluorescent protein cargoes above, initial experiments involved co-infiltration of *N. benthamiana* with *A. tumefaciens* harbouring PDIP fused to P22 SP (SP-PDIP) alongside P22 CP. Although the co-infiltration experiments yielded good CP expression, no SP-PDIP was identified in the ultracentrifuged VLP extracts, suggesting that the SP-PDIP fusion was not packaged into P22 VLPs in *N. benthamiana*. Therefore, an alternative approach of directly fusing the cargo protein to the N-terminus of the CP was employed, as it has been demonstrated to facilitate bioactive peptide encapsulation within P22 VLPs when expressed in *E. coli* (Wang et al., 2018). It was reasoned that direct fusion of PDIP to the N-terminus of the CP (Figure 3A) would still shield PDIP from the plant cell to enable recombinant accumulation if the PDIP-CP fusions assembled into VLPs in *N. benthamiana*.

**Figure 3.**
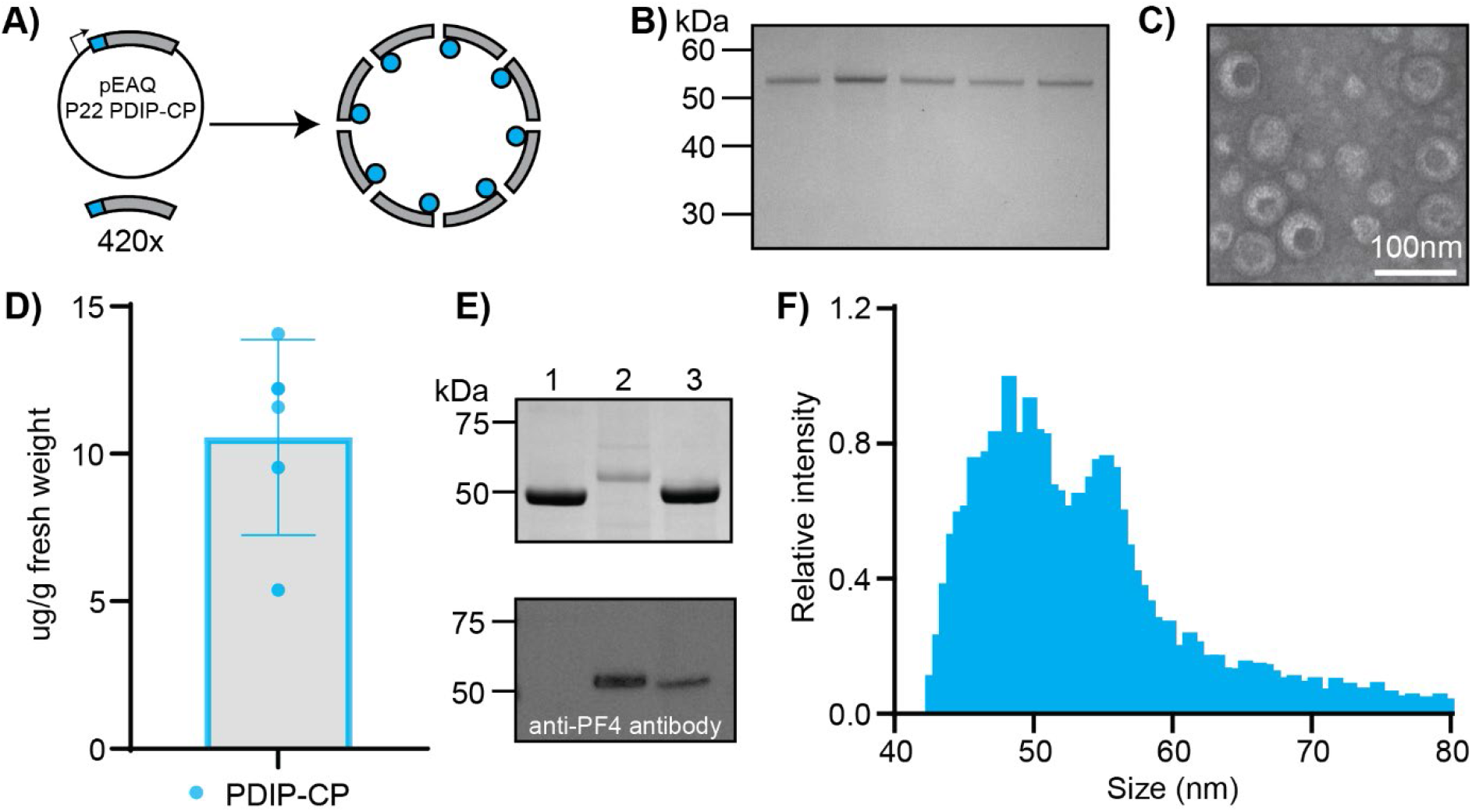
Encapsulation of toxic peptide cargo by fusion to P22 CP. **A)** A cartoon schematic of PDIP-CP fusion construct and its encapsulation within P22 VLPs. **B)** Coomassie stained SDS-PAGE of ultracentrifuged purified protein extracts from transient expressions with PDIP-CP (n=5). **C)** Negative stain TEM of P22 PDIP-CP VLPs. **D)** Calculated yields of PDIP-CP in µg/g fresh weight from n=5 transient expressions. **E)** Coomassie stained SDS-PAGE and corresponding western blot using anti-platelet factor 4 (PF4) antibody, which recognises PDIP. Lanes contain; (1) P22 SP-mRUBY3, (2) P22 PDIP-CP, and (3) chemically synthesised PDIP conjugated to purified P22 VLPs. **F)** Nano flow cytometry size distribution of P22 PDIP-CP VLPs.

Ultracentrifuged protein extracts from transient expressions with PDIP-CP (n=5) were first analysed by SDS-PAGE (Figure 3B). The presence of a single protein species with a mass above 50kDa suggested successful accumulation of PDIP-CP. Subsequent analysis of these samples by negative stain TEM (Figure 3C) confirmed the formation of P22 VLPs. However, heterogenous particle morphology suggests that the N-terminal fusion of PDIP might disrupt CP-CP interactions during VLP assembly. Further analysis of particle morphology by nano flow cytometry revealed a greater spread in size distribution, with a secondary peak shifted closer to 60 nm in diameter compared to the wild-type P22 VLPs loaded with fluorescent proteins (Figure 3F, Figure S3).

To confirm the presence of PDIP in purified PDIP-CP P22 VLPs, a western blot was conducted using an anti-platelet factor 4 polyclonal antibody, which binds to PDIP. P22 SP-mRUBY3 expressed in *N. benthamiana* was used as a negative control (lane 1, Figure 3E). A modified P22 VLP expressed in *E. coli* with chemically synthesised PDIP conjugated to the C-terminus of P22 CP (Harding et al., 2026, unpublished) was used as a positive control (lane 3, Figure 3E). The signal from the western blot for both the positive control and the P22 PDIP-CP VLP sample confirmed the presence of PDIP in the plant produced VLPs. Notably, calculation of the yield of PDIP-CP in µg/g fresh plant weight suggested a significant reduction in CP accumulation compared to the wild-type CP, with an average 10.5 µg/g of PDIP-CP being purified, compared to 196 µg/g of CP purified from the P22 VLPs with SP-mRUBY3 cargo (Figure 3D). The reduced accumulation is consistent with PDIP-mediated toxicity conferred to the host cell during PDIP-CP expression, and that we have previously been unable to purify any recombinantly expressed PDIP from *N. benthamiana*. Regardless of the low recombinant yield, packaging of PDIP into VLPs represents a positive outcome for demonstrating the potential of P22 VLPs to expand the biosynthetic range of *N. benthamiana*.

### Exterior protein display on P22 VLPs using SpyTag/SpyCatcher

A versatile protein scaffold that is amendable to both programmable encapsulation and external display would facilitate a greater diversity of protein engineering applications. To determine whether this could be achieved for P22 VLPs in plants, the ability to functionalise the exterior of P22 VLPs *in vivo* in *N. benthamiana* was next explored. The SpyTag/SpyCatcher (ST/SC) system was employed (Zakeri et al., 2012), as it has been used successfully to functionalise HBc VLPs in the cytosol of *N. benthamiana* (Peyret et al., 2020). The C-terminus of the P22 CP, which is externalised on assembled VLPs, was extended to contain a flexible linker (GSGGGGSG) followed by SpyTag (referred to herein as CP-ST) (Table S1). *N. benthamiana* were co-infiltrated with *A. tumefaciens* strains carrying the modified CP-ST, the SpyCatcher-mCLOVER3 (SC-mCLOVER3) fusion, and SP-mRUBY3, to determine whether *in vivo* particle assembly and ST/SC-mediated external display could be achieved for P22 VLPs produced in *N. benthamiana* (Figure 4A).

**Figure 4.**
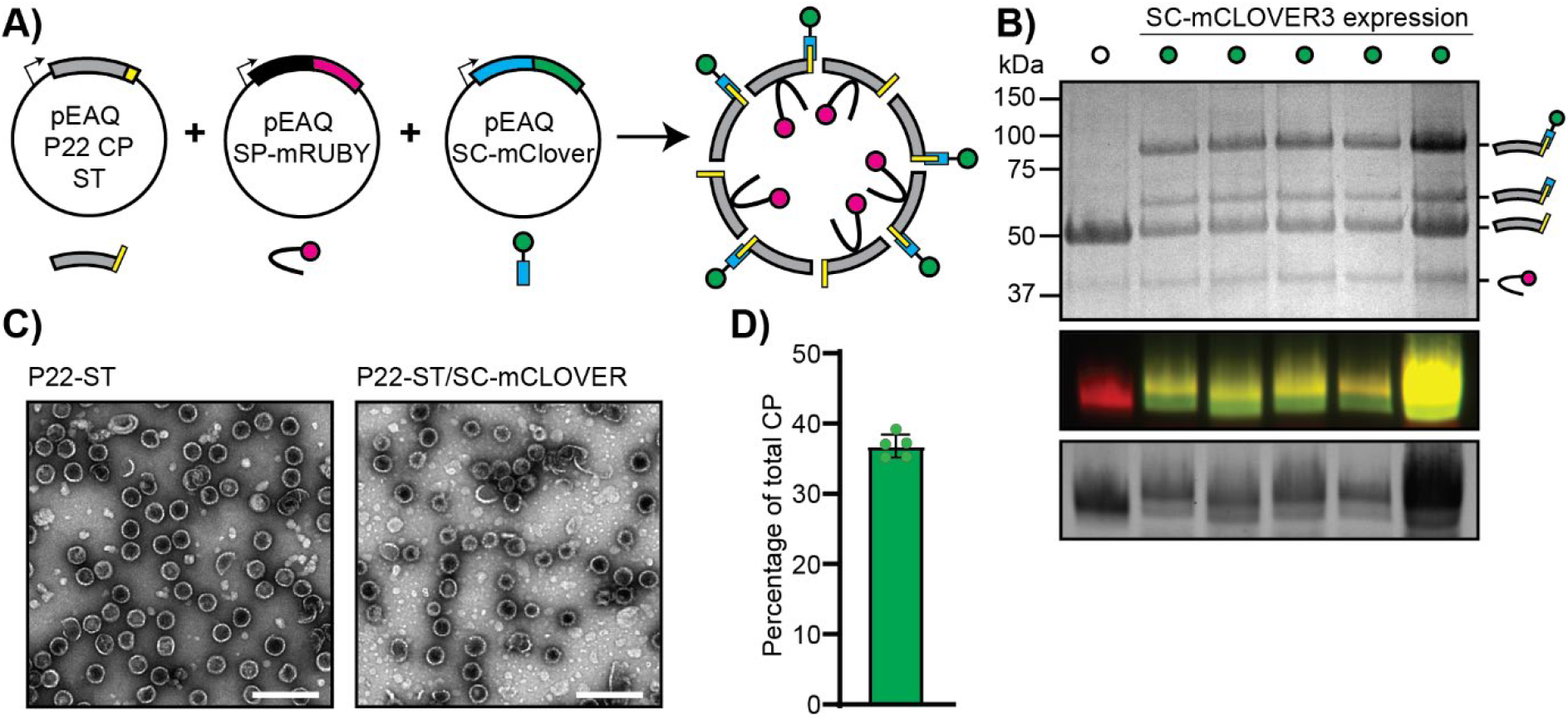
External display of mCLOVER3 on P22 VLPs using SpyTag/SpyCatcher. **A)** Illustrations of the pEAQ vectors harbouring: P22 CP with SpyTag fused to the C-terminus, P22 SP fused to mRUBY3, and SpyCatcher fused to mCLOVER3. The VLP cartoon shows arrangement of the components in the assembled and functionalised P22 VLP. **B)** Coomassie stained gel (SDS-PAGE) of ultracentrifuged protein pellet (top), fluorescent imaging (middle), and a Coomassie stained image of a native agarose gel (bottom). The first lane represents purified P22-ST VLPs without co-expression of SC-mCLOVER3. The remaining five lanes represent purified VLPs with SC-mCLOVER3 co-expressed, as indicated by green dot above lanes (n=5). The protein species of each band on the Coomassie stained gel is indicated by a cartoon illustration on the right. **C)** Negative stain TEM of purified VLPs, labelled above images corresponding to expression without or with SC-mCLOVER3. Scale bar indicates 200nm. **D)** Calculated percentage of SC-mCLOVER3 conjugation as a percentage of the total coat protein.

Initial expression of CP-ST and SP-mRUBY3 produced successfully assembled SpyTagged-VLPs with encapsulated SP-mRUBY3 (Figure 4B lane 1, Figure 4C), confirming that C-terminal extension of the CP with SpyTag does not affect VLP assembly. Co-expression of SC-mCLOVER3 with these constructs produced two higher molecular weight species on SDS-PAGE. The strong band at ∼90 kDa was identified by tryptic digest LC/MS-MS to be full-length CP-ST/SC-mCLOVER3 fusion (expected mass 88.6 kDa, see Table S2). Fluorescence and Coomassie stained imaging of a native agarose gel with these samples also confirmed the presence of both mCLOVER3 and mRUBY3 (Figure 4B), with negative stain TEM demonstrating VLP assembly (Figure 4C) confirming conjugation of SC-mCLOVER3 to the exterior of SpyTagged-VLPs. However, an additional band at ∼60 kDa (Figure 4B) was identified to contain CP-ST/SC, but not mCLOVER3 (Table S2), suggesting that partial cleavage occurred in between SC and mCLOVER3. Densitometry-based quantification revealed an average of 36% of CP-ST successfully conjugated with SC-mCLOVER3 (Figure 4D), which corresponds to approximately 151 SC-mCLOVER3 molecules per SpyTagged-VLP if each particle was correctly assembled with 420 CP-ST monomers. However, given the heterogeneous morphology of VLPs seen by negative stain TEM (Figure 4C), the calculation presented here can only claim to represent the percentage of CP-ST monomers conjugated with SC-mCLOVER3. This level of functionalisation is lower than previously reported for *in vitro* ST/SC-mediated conjugation, which resulted in approximately 230 copies of influenza hemagglutinin head protein per VLP (Sharma et al., 2020) but can be accounted for by the cleavage product which was counted as CP-ST instead of functionalised CP-ST/SC-mCLOVER3.

Formation of ST/SC isopeptide bonds *in vivo* within the same cell is an important requirement for *in planta* applications of externally displayed proteins on P22 VLPs, such as display of enzymes for metabolic applications. Therefore, to evaluate whether SC-mCLOVER3 conjugation to P22 VLPs occurred *in vivo*, or alternatively during cell lysis immediately prior to VLP purification, a plant cell lysate mixing experiment was conducted whereby four plants were co-infiltrated with *A. tumefaciens* harbouring CP-ST and SP-mRUBY3, and another four plants were infiltrated with *A. tumefaciens* harbouring SC-mCLOVER3 alone. Five days post-infiltration, clarified plant protein extracts for SpyTagged-VLPs (CP-ST, SP-mRUBY3) and SC-mCLOVER3 were mixed 1:1 and incubated with shaking at 4 ºC overnight. These samples were then subjected to ultracentrifugation, and the purified VLPs were assessed by SDS-PAGE. In contrast to the co-infiltration experiments described above, VLPs from all four replicate mixing experiments did not show conjugation of SC-mCLOVER3. Additionally, attempts to purify SC-mCLOVER3 from plant extracts via the N-terminal histidine tag using Ni-NTA resin failed to recover any protein, suggesting possible instability of SC-mCLOVER3 alone. Despite the partial cleavage/instability observed for SC-mCLOVER3, these results represent an exciting outcome in demonstrating successful assembly and selective cargo packaging, which can be simultaneously paired with *in vivo* external functionalisation of P22 VLPs in *N. benthamiana*.

### Engineering a protein binding P22 VLP *in planta*

The ability to program both encapsulation of a cargo and external display of a targeting protein on plant-produced P22 VLPs has potential functional applications. The use of VLPs and other protein nanoparticles as targeted drug delivery vehicles is a particularly exciting area of biologics development. Additionally, the formation of higher order P22 VLP assemblies or condensates via external functionalisation has recently been explored for enhancing enzyme catalysis and creating novel biological materials (Zeng et al., 2025, Hewagama et al., 2023). As a proof-of-concept towards producing functional P22 VLPs from plants, we explored fusing three distinct protein binding domains to the SC component of the ST/SC conjugation pair; the anti-GFP nanobody Enhancer which has previously been expressed in plants (Kourelis et al., 2023), and the therapeutic nanobody 9G8 and affibody ZHER2, that target the EGFR and HER2 receptors respectively (Orlova et al., 2006, Roovers et al., 2011) that have not been previously explored for plant expression.

Analysis of purified protein extracts from co-infiltrations with *A. tumefaciens* harbouring CP-ST, SP-mRUBY3, and Enhancer-SC by SDS-PAGE and tryptic digest LC-MS/MS confirmed the conjugation of Enhancer-SC to CP-ST, as seen at ∼70 kDa (expected molecular weight 71.7 kDa, Figure 5B, Table S2), together with a breakage product (∼60 kDa), similar to the outcome for SC-mCLOVER3 co-expression. Negative stain TEM imaging confirmed the formation of VLPs (Figure 5C), with densitometric analysis reporting an average of 37% of total CP-ST functionalised with Enhancer-SC (Figure 5D), in agreement with conjugation efficiency achieved with SC-mCLOVER3.

**Figure 5.**
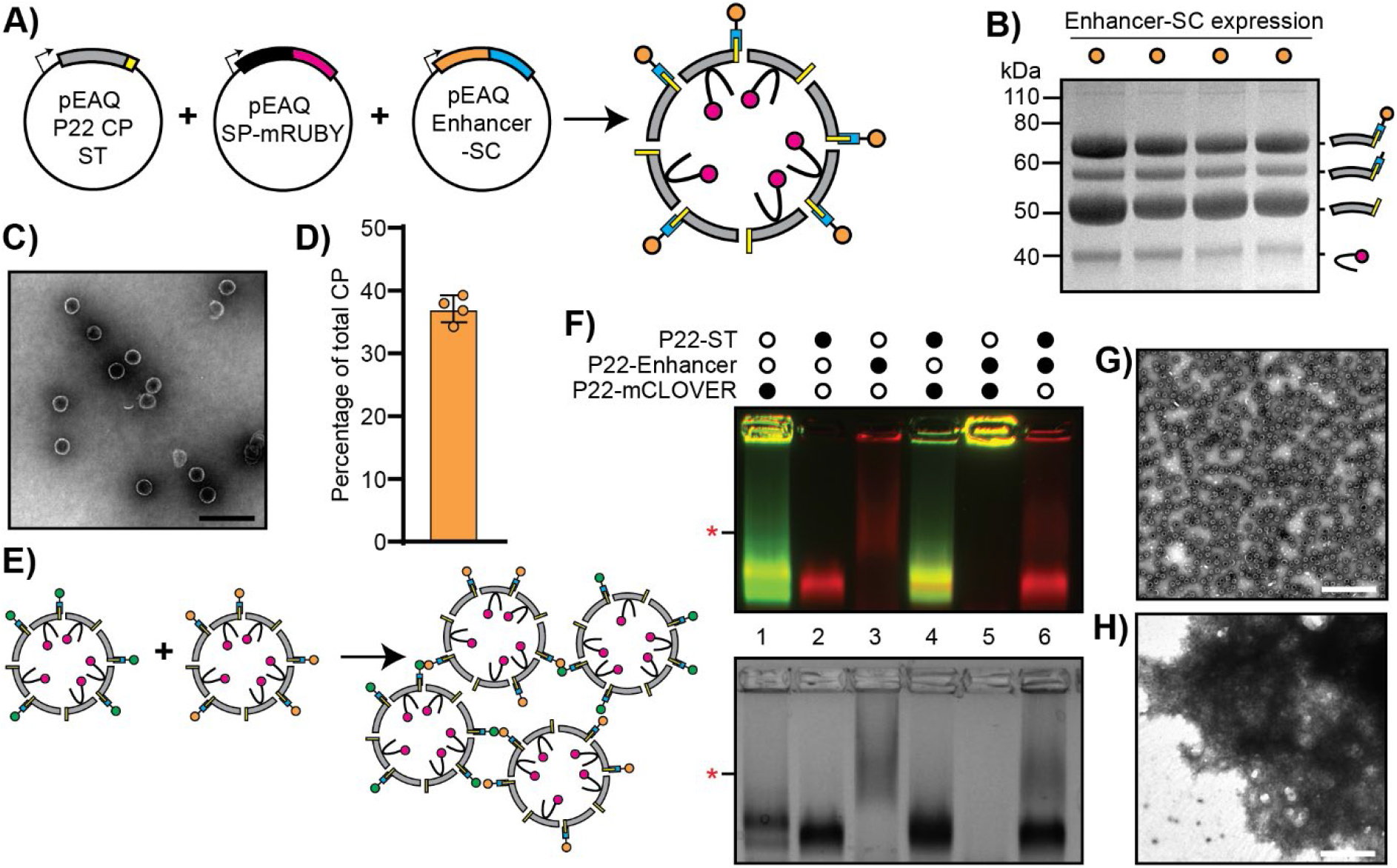
External display of Enhancer nanobody on P22 VLPs using SpyTag/SpyCatche. **A)** Illustrations of the pEAQ vectors harbouring: P22 CP-ST, P22 SP-mRUBY3, and Enhancer-SC fusion. The VLP cartoon shows arrangement of the components in the assembled and functionalised P22 VLP. **B)** Coomassie stained SDS-PAGE of purified VLPs with all three constructs co-expressed (n=4). Protein species indicated by cartoon diagrams on right side of gel. **C)** Negative stain TEM of purified VLPs. Scale bar indicates 200 nm. **D)** Calculated percentage of total CP-ST functionalised with Enhancer-SC (n=4). **E)** Illustration of hypothesised VLP aggregation upon mixing of P22-Enhancer with P22-mCLOVER. **F)** Native agarose gel imaged for fluorescence (top) and stained with Coomassie (bottom). Key at top of gels indicates which VLPs were mixed together prior to loading on gel. Red asterisk indicates where the P22-Enhancer VLP can be seen on the gel in lanes 3 and 6, but not in lane 5. **G)** Negative stain TEM of P22-Enhancer mixed with P22-ST. Scale bar indicates 500 nm. **H)** Negative stain TEM of P22-Enhancer mixed with P22-mCLOVER. Scale bar indicates 500 nm.

To evaluate the functionality of Enhancer-functionalised P22 VLPs, we mixed P22-Enhancer VLPs with P22-mCLOVER VLPs, as we expected that binding between externalised Enhancer and mCLOVER would produce VLP aggregates (Figure 5E). To test this possibility, we conducted an *in vitro* mixing experiment where purified P22-Enhancer VLPs were mixed (1:1) with P22-mCLOVER VLPs, or with non-conjugated VLP-ST (P22-ST) as a control, for 20 min and assessed the outcome by native PAGE (Figure 5F). P22-Enhancer alone ran notably higher on the native PAGE (illustrated by red asterisk, Figure 5F, lane 3) compared to P22-mCLOVER or P22-ST (lane 1 and 2, respectively). This is likely due to lower charge of P22-Enhancer at pH 7.4 (-12 for a single CP-Enhancer fusion) compared to P22-mCLOVER (-21 for a single CP-mCLOVER fusion) and a greater size compared to non-functionalised P22-ST (-9 per CP-ST). Control incubations with P22-ST resulted in no change to the native PAGE signal when mixed with P22-mCLOVER (lane 4) and P22-Enhancer (lane 6). However, mixing of P22-Enhancer with P22-mCLOVER restricted fluorescence to the top of the well (lane 5) indicating VLP aggregation. This VLP aggregate signal could not be seen upon Coomassie staining, possibly due to the protein washing away upon destaining given its inability to pass into the gel matrix. Assessment of P22-Enhancer mixed with P22-ST by negative stain TEM displayed clear individual VLPs (Figure 5G). However, the P22-Enhancer mixed with P22-mCLOVER showed extensive aggregation, as seen in the TEM micrograph (Figure 5H). These results suggest that the Enhancer nanobody is functional when conjugated to the exterior of P22 VLPs via SpyTag/SpyCatcher in plants. Furthermore, the data suggest that exterior display of nanobodies on P22 VLPs can be used to create higher-order VLP assemblies *in vitro*.

In contrast to the expression and functionalisation with Enhancer, *in planta* expression of 9G8-SC nanobody or SC-ZHER2 affibody did not produce functionalised VLPs under the conditions tested. Although the 9G8-SC nanobody could be expressed by *N. benthamiana*, VLPs functionalised with 9G8-SC were insoluble in PBS (Figure S4). Interestingly, a fraction of VLPs which had 9G8 cleaved were soluble (Figure S4), demonstrating that 9G8-SC fusion and display impacts VLP solubility. Similarly, functionalisation with SC-ZHER2 was not readily achieved as only P22 VLPs with the SC cleavage product could be purified from *N. benthamiana* expressions (Figure S5). However, SC-ZHER2 could be expressed in *E. coli* and successfully conjugated to plant expressed VLP-ST *in vitro* (Figure S5), suggesting that the ZHER2 affibody is unstable in *N. benthamiana*. The recurrence of a cleavage product and reduced solubility for certain constructs suggests that further optimisation – either at the level of construct design or during downstream processing – may be required to accommodate a broader range of functional external cargoes. Collectively, the results for mCLOVER3, Enhancer, and the two therapeutic protein binders 9G8 and ZHER2 demonstrate the ability to program protein display on the exterior of P22 VLPs with ST/SC *in vivo*.

## Conclusion

The biofactory plant *N. benthamiana* is a flexible host for plant synthetic biology, with new tools for recombinant production of high-value molecules continuing to be developed. Here, we explored the use of bacteriophage P22 VLPs as a new protein scaffold for synthetic biology applications in *N. benthamiana*. Through co-expression of the P22 CP with proteins of interest fused to the P22 SP, the selective encapsulation of protein cargo within P22 VLPs can be achieved (Figure 6A). This success extends to the co-encapsulation of at least two different proteins – as demonstrated with mRUBY3 and mCLOVER3 – but could be expanded to include a greater number of different cargoes (Figure 6B), such as has been demonstrated for metabolic engineering applications in *E. coli* (McNeale et al., 2023b). Through genetic fusion to either the CP N-terminus (with PDIP for encapsulation) or C-terminus (with SpyTag for external display), we show that P22 VLPs maintain their robust assembly capacity in plants (Figure 6C, 6D). Via co-expression of SpyCatcher-mCLOVER3 fusions with CP-SpyTag, we show that P22 VLPs can be externally conjugated with proteins of interest *in vivo* – achieving two layers of programmability; cargo encapsulation and external protein display. We further expand this capability by creating functional VLPs displaying the anti-GFP nanobody Enhancer that bind P22 displaying mCLOVER3 *in vitro* to form higher-order VLP assemblies.

**Figure 6.**
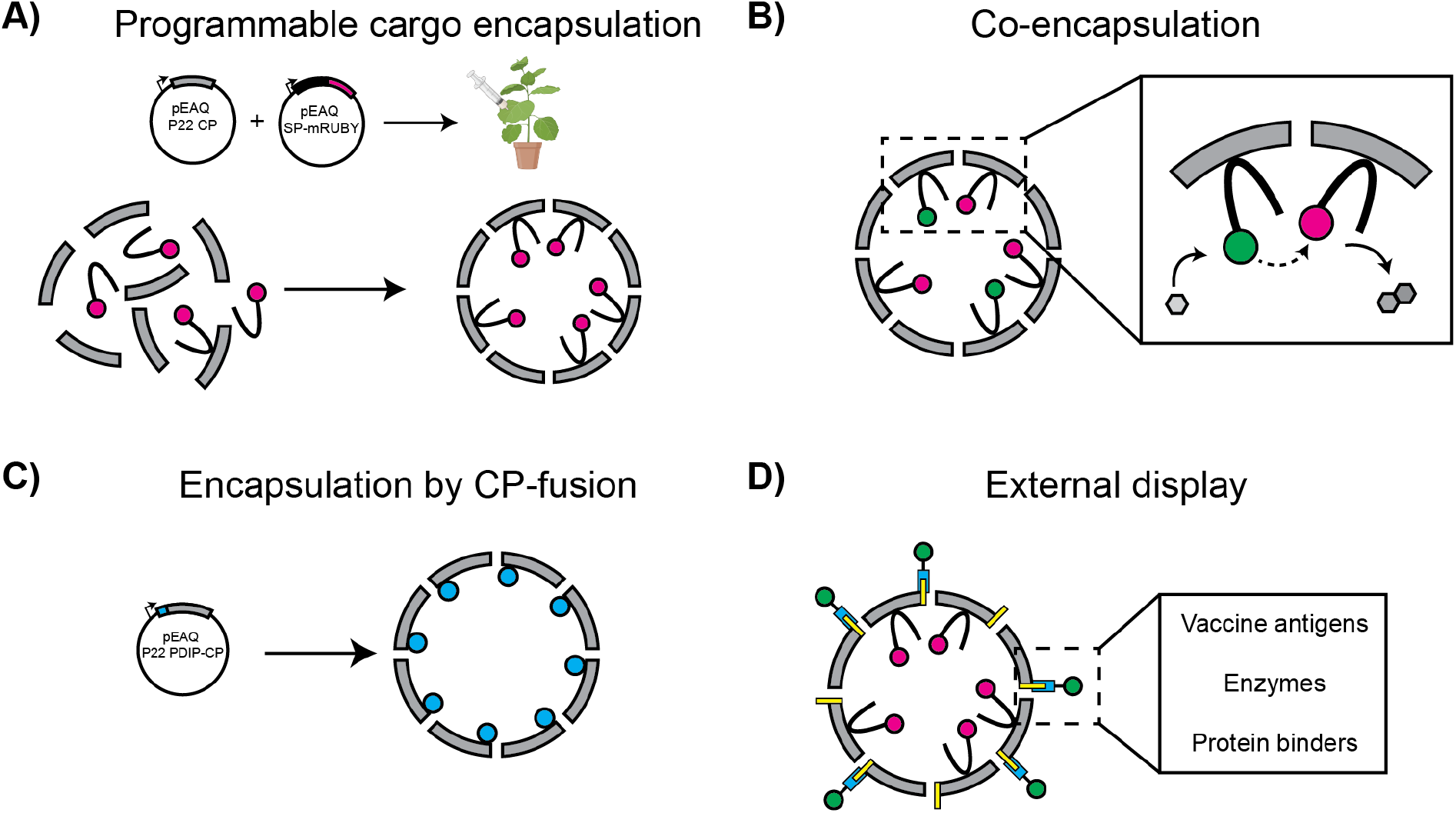
Applications of P22 VLPs as scaffolds for protein engineering in *N. benthamiana*. These applications include (**A**) programmable cargo encapsulation for both a single protein cargo; (**B**) co-encapsulation of two protein cargoes; (**C)** encapsulation by fusion directly to the N-terminus of the P22 CP; and (**D)** external display of proteins mediated by SpyTag/SpyCatcher.

In summary, the work presented here demonstrates the utility of P22 VLPs as highly programmable protein scaffolds, capable of both encapsulation and display of engineered proteins in *N. benthamiana*. However, there are some limitations of the system. Cargo encapsulation by SP fusion or external display by fusion to SC need to be evaluated on a case-by-case basis, as is the case for most recombinant proteins expressed in plants. Specifically for external functionalisation, all SC-fused proteins evaluated here displayed a breakage product which may be occurring around the flexible linker sequence between SC and the fusion partner, or within SC itself. Overall, the amenability of P22 VLPs to protein engineering makes this scaffold poised for novel applications in plant synthetic biology. Programming *in vivo* encapsulation could be expanded to therapeutic cargo proteins or other biomolecules such as nucleic acid, with applications ranging from metabolic engineering to the development of delivery vehicles for biomedicine or agriculture, and possibly to synthetic biology in other eukaryotes.

## Methodology

### Cloning

Proteins for transient expression were ordered as gene blocks from Integrated DNA Technologies in the form of dsDNA. These were cloned using Gateway cloning enzymes (BP and LR Clonase II from Invitrogen) first into the pDS221 intermediate vector and then into the plant expression vector pEAQ-DEST1 (Sainsbury et al., 2009). All plasmids were confirmed by sequencing using either Sanger sequencing (Australian Genome Research Facility) or by whole plasmid long read sequencing (Plasmidsaurus). Sequenced plasmids were then electroporated into *A. tumefaciens* strain LBA4404 for transient expression.

### Transient expression in *N. benthamiana*

Transformed *A. tumefaciens* were grown in Luria-Bertani medium supplemented with 50 µg/mL kanamycin and pelleted by centrifugation at 3,900 g. Cells were then resuspended in infiltration buffer (10 mM MES (2-[*N*-morpholino]ethanesulfonic acid), 10mM MgSO_4_, 100 µM acetosyringone, pH 5.6) and left to rest for 2 h at room temperature. Different *A. tumefaciens* strains co-infiltrated together were mixed at equal amounts to ensure a final A_600_ of 1.0 by dilution with infiltration buffer. *N. benthamiana* plants between 4 to 6 weeks old were then transiently transformed by syringe infiltration to the abaxial side of the leaf. Infiltrated plant tissue was harvested 5 days post-infiltration and fresh weight recorded.

### Protein purification and analysis

Leaf tissue was ground in liquid nitrogen before incubation on ice in VLP extraction buffer (50 mM bicine, 140 mM NaCl, 1 mM DTT, 0.1% w/v NLS sodium salt, EDTA-free protease inhibitor, pH 8.4). Plant protein extracts were filtered through miracloth, then centrifuged at 20,000 *g* for 20 min at 4 ºC. Clarified extract was then filtered (0.45 µm) before being applied to an iodixanol gradient (15, 20, 25%) in extraction buffer and ultracentrifuged for 3 h at 150,000 g. The bottom pellet of the ultracentrifuge tube was extracted and desalted into PBS using a PD10-miditrap column (Cytiva). Purified protein was quantified by a Pierce BCA assay (ThermoFisher Scientific) and 5 µg protein for each sample analysed by denaturing SDS-PAGE and Coomassie staining. Cargo protein loading was calculated by densitometry using ImageJ software, assuming correct VLP assembly with 420 CP monomers per VLP. Protein species were identified by in-gel tryptic digest liquid LC-MS/MS on a Sciex 5600 Quadrupole time-of-flight mass spectrometer (QTOF-MS). Western blot analysis of PDIP-CP was conducted using a rabbit polyclonal anti-PF4 primary (ABCAM Australia) and a horseradish peroxidase-fused anti-rabbit secondary antibody. The blot was visualised on a ChemiDoc MP (BioRad, Australia) following application of SuperSignal West Pico PLUS chemiluminescent substrate (ThermoFisher). NAGE was conducted by running 10 µg of purified VLP on a 0.8% agarose gel. For analysis of nucleic acid packaging, gels were stained post-electrophoresis with SYBR Safe for 30 min before destaining and imaging. Destained gels were then stained over night with Coomassie. Fluorescent images were taken on a BioRad ChemiDoc imaging system using GFP (green) and mCherry (red) channels.

### Negative stain transmission electron microscopy

Purified VLPs (10 µL, 0.25 mg/mL) were applied to formvar/carbon-coated copper grids (ProSciTech) for 2 min. Grids were then washed three times in 10 µL water before staining in 2% uranyl acetate for 1 min. Imagining was performed on a JEOL 1400 Flash electron microscope at 80 kV operating voltage.

### Förster resonance energy transfer experiment

Each sample was normalised to 20 µg/mL of P22 CP in PBS with three technical replicates (50 µL) of each analysed in a black 96-well microtiter plate. Samples were excited at 450 nm and emissions recorded at 518 nm. Emission spectra were averaged from the three technical replicates, had the PBS baseline subtracted and were normalised to the emission maximum at 518 nm.

### Nanoflow cytometry

Samples were analysed on a U30E NanoAnalyser (NanoFCM), with a dilution series starting from 0.1 mg/mL.

### *E. coli* protein expression and purification

BL21(DE3) cells transformed with pET20b(+) containing SC-ZHER2 were induced at an A_600_ of 0.6 and left overnight at 18 ºC. Cells were harvested by centrifugation at 4,500 *g* for 15 min. Lysed cells were then clarified at 40,000 *g* for 30 min, before being passed through a 0.45 µm filter and loaded onto a 5 mL HisTrap FF column (Cytiva) and purified using an ÄKTA FPLC system. Fractions containing SC-ZHER2 were concentrated and buffer exchanged into PBS pH 7.4 using a 3.5 kDa Amicon filter before being quantified by A_280_ absorbance using a NanoDrop.

## Author contributions

**Maxim D. Harding** conceptualization, data curation, formal analysis, investigation, methodology, project administration, validation, visualization, writing – original draft; **Mark A. Jackson** funding acquisition, methodology, resources, supervision, writing – review and editing; **Edward K. Gilding** funding acquisition, methodology, supervision, writing – review and editing; **David J. Craik** funding acquisition, resources, supervision, writing – review and editing; **Frank Sainsbury** conceptualization, methodology, resources, validation, writing – review and editing; **Nicole Lawrence** funding acquisition, methodology, project administration, resources, supervision, validation, writing – review and editing.

## Acknowledgements

This work was supported by funding from the US Department of Defense grant (PR210354 to DJC, NL, MAJ, EKG) and the Australian Research Council Centre of Excellence for Innovations in Peptide and Protein Science (CE200100012). DJC was supported by NHMRC grant (2009564). FS is supported by an Australian Research Council Future Fellowship (FT230100084) and the Australian Research Council Centre of Excellence in Synthetic Biology (CE200100029).

## Data availability statement

All data is presented in the manuscript text and/or supporting information.

