## Supplementary Information for "Bacteriophage P22 virus-like particles as nanoscale protein scaffolds for plant synthetic biology"

**Contents**

Table S1. Amino acid sequence of proteins used in this study

Table S2. Tryptic digest LC-MS/MS of recombinant proteins

Table S3. Nucleic acid sequence of CP ST_LS1

Figure S1. Intron-containing CP expression

Figure S2. Fluorescent imaging of infiltrated leaf tissue

Figure S3. Nano flow cytometry of purified VLPs

Figure S4. Exploring display of 9G8 nanobody

Figure S5. Exploring display of ZHER2 affibody

**Table S1. Amino acid sequence of proteins used in this study**

| **Protein** | **Sequence** | **MW (kDa)** |
| --- | --- | --- |
| Coat Protein (CP) | MALNEGQIVTLAVDEIIETISAITPMAQKAKKYTPPAASMQRSSNTIWMPVEQ  ESPTQEGWDLTDKATGLLELNVAVNMGEPDNDFFQLRADDLRDETAYRRRIQS  AARKLANNVELKVANMAAEMGSLVITSPDAIGTNTADAWNFVADAEEIMFSRE  LNRDMGTSYFFNPQDYKKAGYDLTKRDIFGRIPEEAYRDGTIQRQVAGFDDVL  RSPKLPVLTKSTATGITVSGAQSFKPVAWQLDNDGNKVNVDNRFATVTLSATT  GMKRGDKISFAGVKFLGQMAKNVLAQDATFSVVRVVDGTHVEITPKPVALDDV  SLSPEQRAYANVNTSLADAMAVNILNVKDARTNVFWADDAIRIVSQPIPANHE  LFAGMKTTSFSIPDVGLNGIFATQGDISTLSGLCRIALWYGVNATRPEAIGVG  LPGQTA | 46.75 |
| Scaffold protein (SP)  SP-mRUBY3 | MTRLSERLTLKPRGKQISSAPPADQPITGDVSAANKDAIRKQMDAAASKGDVE  TYRKLKAKLKGIRGGGGSGGGGSMVSKGEELIKENMRMKVVMEGSVNGHQFKC  TGEGEGRPYEGVQTMRIKVIEGGPLPFAFDILATSFMYGSRTFIKYPADIPDF  FKQSFPEGFTWERVTRYEDGGVVTVTQDTSLEDGELVYNVKVRGVNFPSNGPV  MQKKTKGWEPNTEMMYPADGGLRGYTDIALKVDGGGHLHCNFVTTYRSKKTVG  NIKMPGVHAVDHRLERIEESDNETYVVQREVAVAKYSNLGGGMDELYKGSHHH  HHH | 35.32 |
| SP-mCLOVER3 | MTRLSERLTLKPRGKQISSAPPADQPITGDVSAANKDAIRKQMDAAASKGDVE  TYRKLKAKLKGIRGGGGSGGGGMVSKGEELFTGVVPILVELDGDVNGHKFSVR  GEGEGDATNGKLTLKFICTTGKLPVPWPTLVTTFGYGVACFSRYPDHMKQHDF  FKSAMPEGYVQERTISFKDDGTYKTRAEVKFEGDTLVNRIELKGIDFKEDGNI  LGHKLEYNFNSHYVYITADKQKNCIKANFKIRHNVEDGSVQLADHYQQNTPIG  DGPVLLPDNHYLSHQSKLSKDPNEKRDHMVLLEFVTAAGITHGMDELYK | 34.65 |
| PDIP-CP | MGCGAPLYKKIIKKLLESGGSGGAPLYKKIIKKLCESNGLMALNEGQIVTLAV  DEIIETISAITPMAQKAKKYTPPAASMQRSSNTIWMPVEQESPTQEGWDLTDK  ATGLLELNVAVNMGEPDNDFFQLRADDLRDETAYRRRIQSAARKLANNVELKV  ANMAAEMGSLVITSPDAIGTNTADAWNFVADAEEIMFSRELNRDMGTSYFFNP  QDYKKAGYDLTKRDIFGRIPEEAYRDGTIQRQVAGFDDVLRSPKLPVLTKSTA  TGITVSGAQSFKPVAWQLDNDGNKVNVDNRFATVTLSATTGMKRGDKISFAGV  KFLGQMAKNVLAQDATFSVVRVVDGTHVEITPKPVALDDVSLSPEQRAYANVN  TSLADAMAVNILNVKDARTNVFWADDAIRIVSQPIPANHELFAGMKTTSFSIP  DVGLNGIFATQGDISTLSGLCRIALWYGVNATRPEAIGVGLPGQTA | 50.94 |
| CP-SpyTag  (CP-ST) | MALNEGQIVTLAVDEIIETISAITPMAQKAKKYTPPAASMQRSSNTIWMPVEQ  ESPTQEGWDLTDKATGLLELNVAVNMGEPDNDFFQLRADDLRDETAYRRRIQS  AARKLANNVELKVANMAAEMGSLVITSPDAIGTNTADAWNFVADAEEIMFSRE  LNRDMGTSYFFNPQDYKKAGYDLTKRDIFGRIPEEAYRDGTIQRQVAGFDDVL  RSPKLPVLTKSTATGITVSGAQSFKPVAWQLDNDGNKVNVDNRFATVTLSATT  GMKRGDKISFAGVKFLGQMAKNVLAQDATFSVVRVVDGTHVEITPKPVALDDV  SLSPEQRAYANVNTSLADAMAVNILNVKDARTNVFWADDAIRIVSQPIPANHE  LFAGMKTTSFSIPDVGLNGIFATQGDISTLSGLCRIALWYGVNATRPEAIGVG  LPGQTAGSGGGGSGAHIVMVDAYKPTK | 48.72 |
| SpyCatcher-mCLOVER3  (SC-mCLOVER3) | MSYYHHHHHHDYDIPTTENLYFQGSGDSATHIKFSKRDEDGKELAGATMELRD  SSGKTISTWISDGQVKDFYLYPGKYTFVETAAPDGYEVATAITFTVNEQGQVT  VNGKATKGSGGGSGGGMVSKGEELFTGVVPILVELDGDVNGHKFSVRGEGEGD  ATNGKLTLKFICTTGKLPVPWPTLVTTFGYGVACFSRYPDHMKQHDFFKSAMP  EGYVQERTISFKDDGTYKTRAEVKFEGDTLVNRIELKGIDFKEDGNILGHKLE  YNFNSHYVYITADKQKNCIKANFKIRHNVEDGSVQLADHYQQNTPIGDGPVLL  PDNHYLSHQSKLSKDPNEKRDHMVLLEFVTAAGITHGMDELYK | 40.15 |

| Enhancer-SpyCatcher  (Enhancer-SC) | MAQVQLVESGGALVQPGGSLRLSCAASGFPVNRYSMRWYRQAPGKEREWVAGM  SSAGDRSSYEDSVKGRFTISRDDARNTVYLQMNSLKPEDTAVYYCNVNVGFEY  WGQGTQVTVSSSSGGGSGGGDSATHIKFSKRDEDGKELAGATMELRDSSGKTI  STWISDGQVKDFYLYPGKYTFVETAAPDGYEVATAITFTVNEQGQVTVNGKAT  KG | 23.05 |
| --- | --- | --- |
| SpyCatcher-ZHER2  (SC-ZHER2) | MSYYHHHHHHDYDIPTTENLYFQGSGDSATHIKFSKRDEDGKELAGATMELRD  SSGKTISTWISDGQVKDFYLYPGKYTFVETAAPDGYEVATAITFTVNEQGQVT  VNGKATKGSGGGSGGGVDNKFNKEMRNAYWEIALLPNLNNQQKRAFIRSLYDD  PSQSANLLAEAKKLNDAQAPK | 19.92 |
| 9G8-SpyCatcher  (9G8-SC) | MEVQLVESGGGLVQAGGSLRLSCAASGRTFSSYAMGWFRQAPGKEREFVVAIN  WSSGSTYYADSVKGRFTISRDNAKNTMYLQMNSLKPEDTAVYYCAAGYQINSG  NYNFKDYEYDYWGQGTQVTVSSSGGGSGGGDSATHIKFSKRDEDGKELAGATM  ELRDSSGKTISTWISDGQVKDFYLYPGKYTFVETAAPDGYEVATAITFTVNEQ  GQVTVNGKATKG | 24.20 |

**Table S2. Tryptic digest LC-MS/MS of recombinant proteins**

| **Protein** | **Sequence^a^** |
| --- | --- |
| CP | MALNEGQIVTLAVDEIIETISAITPMAQKAKKYTPPAASMQRSSNTIWMPVEQESPTQEGWDLTD  KATGLLELNVAVNMGEPDNDFFQLRADDLRDETAYRRRIQSAARKLANNVELKVANMAAEMGSLV  ITSPDAIGTNTADAWNFVADAEEIMFSRELNRDMGTSYFFNPQDYKKAGYDLTKRDIFGRIPEEA  YRDGTIQRQVAGFDDVLRSPKLPVLTKSTATGITVSGAQSFKPVAWQLDNDGNKVNVDNRFATVT  LSATTGMKRGDKISFAGVKFLGQMAKNVLAQDATFSVVRVVDGTHVEITPKPVALDDVSLSPEQR  AYANVNTSLADAMAVNILNVKDARTNVFWADDAIRIVSQPIPANHELFAGMKTTSFSIPDVGLNG  IFATQGDISTLSGLCRIALWYGVNATRPEAIGVGLPGQTA |
| SP-mRUBY3 | MTRLSERLTLKPRGKQISSAPPADQPITGDVSAANKDAIRKQMDAAASKGDVETYRKLKAKLKGI  RGGGGSGGGGSMVSKGEELIKENMRMKVVMEGSVNGHQFKCTGEGEGRPYEGVQTMRIKVIEGGP  LPFAFDILATSFMYGSRTFIKYPADIPDFFKQSFPEGFTWERVTRYEDGGVVTVTQDTSLEDGEL  VYNVKVRGVNFPSNGPVMQKKTKGWEPNTEMMYPADGGLRGYTDIALKVDGGGHLHCNFVTTYRS  KKTVGNIKMPGVHAVDHRLERIEESDNETYVVQREVAVAKYSNLGGGMDELYKGSHHHHHH |
| SP-mCLOVER3 | MTRLSERLTLKPRGKQISSAPPADQPITGDVSAANKDAIRKQMDAAASKGDVETYRKLKAKLKGI  RGGGGSGGGGMVSKGEELFTGVVPILVELDGDVNGHKFSVRGEGEGDATNGKLTLKFICTTGKLP  VPWPTLVTTFGYGVACFSRYPDHMKQHDFFKSAMPEGYVQERTISFKDDGTYKTRAEVKFEGDTL  VNRIELKGIDFKEDGNILGHKLEYNFNSHYVYITADKQKNCIKANFKIRHNVEDGSVQLADHYQQ  NTPIGDGPVLLPDNHYLSHQSKLSKDPNEKRDHMVLLEFVTAAGITHGMDELYK |
| CP-ST_LS1 | MALNEGQIVTLAVDEIIETISAITPMAQKAKKYTPPAASMQRSSNTIWMPVEQESPTQEGWDLTD  KATGLLELNVAVNMGEPDNDFFQLRADDLRDETAYRRRIQSAARKLANNVELKVANMAAEMGSLV  ITSPDAIGTNTADAWNFVADAEEIMFSRELNRDMGTSYFFNPQDYKKAGYDLTKRDIFGRIPEEA  YRDGTIQRQVAGFDDVLRSPKLPVLTKSTATGITVSGAQSFKPVAWQLDNDGNKVNVDNRFATVT  LSATTGMKRGDKISFAGVKFLGQMAKNVLAQDATFSVVRVVDGTHVEITPKPVALDDVSLSPEQR  AYANVNTSLADAMAVNILNVKDARTNVFWADDAIRIVSQPIPANHELFAGMKTTSFSIPDVGLNG  IFATQGDISTLSGLCRIALWYGVNATRPEAIGVGLPGQTA |
| CP-SpyTag | MALNEGQIVTLAVDEIIETISAITPMAQKAKKYTPPAASMQRSSNTIWMPVEQESPTQEGWDLTD  KATGLLELNVAVNMGEPDNDFFQLRADDLRDETAYRRRIQSAARKLANNVELKVANMAAEMGSLV  ITSPDAIGTNTADAWNFVADAEEIMFSRELNRDMGTSYFFNPQDYKKAGYDLTKRDIFGRIPEEA  YRDGTIQRQVAGFDDVLRSPKLPVLTKSTATGITVSGAQSFKPVAWQLDNDGNKVNVDNRFATVT  LSATTGMKRGDKISFAGVKFLGQMAKNVLAQDATFSVVRVVDGTHVEITPKPVALDDVSLSPEQR  AYANVNTSLADAMAVNILNVKDARTNVFWADDAIRIVSQPIPANHELFAGMKTTSFSIPDVGLNG  IFATQGDISTLSGLCRIALWYGVNATRPEAIGVGLPGQTAGSGGGGSGAHIVMVDAYKPTK |
| CP-SpyTag-SpyCatcher-mCLOVER3 | MALNEGQIVTLAVDEIIETISAITPMAQKAKKYTPPAASMQRSSNTIWMPVEQESPTQEGWDLTD  KATGLLELNVAVNMGEPDNDFFQLRADDLRDETAYRRRIQSAARKLANNVELKVANMAAEMGSLV  ITSPDAIGTNTADAWNFVADAEEIMFSRELNRDMGTSYFFNPQDYKKAGYDLTKRDIFGRIPEEA  YRDGTIQRQVAGFDDVLRSPKLPVLTKSTATGITVSGAQSFKPVAWQLDNDGNKVNVDNRFATVT  LSATTGMKRGDKISFAGVKFLGQMAKNVLAQDATFSVVRVVDGTHVEITPKPVALDDVSLSPEQR  AYANVNTSLADAMAVNILNVKDARTNVFWADDAIRIVSQPIPANHELFAGMKTTSFSIPDVGLNG  IFATQGDISTLSGLCRIALWYGVNATRPEAIGVGLPGQTAGSGGGGSGAHIVMVDAYKPTKMSYY  HHHHHHDYDIPTTENLYFQGSGDSATHIKFSKRDEDGKELAGATMELRDSSGKTISTWISDGQVK  DFYLYPGKYTFVETAAPDGYEVATAITFTVNEQGQVTVNGKATKGSGGGSGGGMVSKGEELFTGV  VPILVELDGDVNGHKFSVRGEGEGDATNGKLTLKFICTTGKLPVPWPTLVTTFGYGVACFSRYPD  HMKQHDFFKSAMPEGYVQERTISFKDDGTYKTRAEVKFEGDTLVNRIELKGIDFKEDGNILGHKL  EYNFNSHYVYITADKQKNCIKANFKIRHNVEDGSVQLADHYQQNTPIGDGPVLLPDNHYLSHQSK  LSKDPNEKRDHMVLLEFVTAAGITHGMDELYK |
| CP-SpyTag-SpyCatcher breakage (~60 kDa) for CP-ST/SC-mCLOVER3 | MALNEGQIVTLAVDEIIETISAITPMAQKAKKYTPPAASMQRSSNTIWMPVEQESPTQEGWDLTD  KATGLLELNVAVNMGEPDNDFFQLRADDLRDETAYRRRIQSAARKLANNVELKVANMAAEMGSLV  ITSPDAIGTNTADAWNFVADAEEIMFSRELNRDMGTSYFFNPQDYKKAGYDLTKRDIFGRIPEEA  YRDGTIQRQVAGFDDVLRSPKLPVLTKSTATGITVSGAQSFKPVAWQLDNDGNKVNVDNRFATVT  LSATTGMKRGDKISFAGVKFLGQMAKNVLAQDATFSVVRVVDGTHVEITPKPVALDDVSLSPEQR  AYANVNTSLADAMAVNILNVKDARTNVFWADDAIRIVSQPIPANHELFAGMKTTSFSIPDVGLNG  IFATQGDISTLSGLCRIALWYGVNATRPEAIGVGLPGQTAGSGGGGSGAHIVMVDAYKPTKMSYY  HHHHHHDYDIPTTENLYFQGSGDSATHIKFSKRDEDGKELAGATMELRDSSGKTISTWISDGQVK  DFYLYPGKYTFVETAAPDGYEVATAITFTVNEQGQVTVNGKATKGSGGGSGGGMVSKGEELFTGV  VPILVELDGDVNGHKFSVRGEGEGDATNGKLTLKFICTTGKLPVPWPTLVTTFGYGVACFSRYPD  HMKQHDFFKSAMPEGYVQERTISFKDDGTYKTRAEVKFEGDTLVNRIELKGIDFKEDGNILGHKL  EYNFNSHYVYITADKQKNCIKANFKIRHNVEDGSVQLADHYQQNTPIGDGPVLLPDNHYLSHQSK  LSKDPNEKRDHMVLLEFVTAAGITHGMDELYK |
| CP-ST/Enhancer-SC | MALNEGQIVTLAVDEIIETISAITPMAQKAKKYTPPAASMQRSSNTIWMPVEQESPTQEGWDLTD  KATGLLELNVAVNMGEPDNDFFQLRADDLRDETAYRRRIQSAARKLANNVELKVANMAAEMGSLV  ITSPDAIGTNTADAWNFVADAEEIMFSRELNRDMGTSYFFNPQDYKKAGYDLTKRDIFGRIPEEA  YRDGTIQRQVAGFDDVLRSPKLPVLTKSTATGITVSGAQSFKPVAWQLDNDGNKVNVDNRFATVT  LSATTGMKRGDKISFAGVKFLGQMAKNVLAQDATFSVVRVVDGTHVEITPKPVALDDVSLSPEQR  AYANVNTSLADAMAVNILNVKDARTNVFWADDAIRIVSQPIPANHELFAGMKTTSFSIPDVGLNG  IFATQGDISTLSGLCRIALWYGVNATRPEAIGVGLPGQTAGSGGGGSGAHIVMVDAYKPTKMAQV  QLVESGGALVQPGGSLRLSCAASGFPVNRYSMRWYRQAPGKEREWVAGMSSAGDRSSYEDSVKGR  FTISRDDARNTVYLQMNSLKPEDTAVYYCNVNVGFEYWGQGTQVTVSSSSGGGSGGGDSATHIKF  SKRDEDGKELAGATMELRDSSGKTISTWISDGQVKDFYLYPGKYTFVETAAPDGYEVATAITFTV  NEQGQVTVNGKATKG |
| CP-ST/SC-ZHER2 (*in vitro* conjugation from *E. coli* expression) | MALNEGQIVTLAVDEIIETISAITPMAQKAKKYTPPAASMQRSSNTIWMPVEQESPTQEGWDLTD  KATGLLELNVAVNMGEPDNDFFQLRADDLRDETAYRRRIQSAARKLANNVELKVANMAAEMGSLV  ITSPDAIGTNTADAWNFVADAEEIMFSRELNRDMGTSYFFNPQDYKKAGYDLTKRDIFGRIPEEA  YRDGTIQRQVAGFDDVLRSPKLPVLTKSTATGITVSGAQSFKPVAWQLDNDGNKVNVDNRFATVT  LSATTGMKRGDKISFAGVKFLGQMAKNVLAQDATFSVVRVVDGTHVEITPKPVALDDVSLSPEQR  AYANVNTSLADAMAVNILNVKDARTNVFWADDAIRIVSQPIPANHELFAGMKTTSFSIPDVGLNG  IFATQGDISTLSGLCRIALWYGVNATRPEAIGVGLPGQTAGSGGGGSGAHIVMVDAYKPTKMSYY  HHHHHHDYDIPTTENLYFQGSGDSATHIKFSKRDEDGKELAGATMELRDSSGKTISTWISDGQVK  DFYLYPGKYTFVETAAPDGYEVATAITFTVNEQGQVTVNGKATKGSGGGSGGGVDNKFNKEMRNA  YWEIALLPNLNNQQKRAFIRSLYDDPSQSANLLAEAKKLNDAQAPK |
| CP-ST/9G8-SC | MALNEGQIVTLAVDEIIETISAITPMAQKAKKYTPPAASMQRSSNTIWMPVEQESPTQEGWDLTD  KATGLLELNVAVNMGEPDNDFFQLRADDLRDETAYRRRIQSAARKLANNVELKVANMAAEMGSLV  ITSPDAIGTNTADAWNFVADAEEIMFSRELNRDMGTSYFFNPQDYKKAGYDLTKRDIFGRIPEEA  YRDGTIQRQVAGFDDVLRSPKLPVLTKSTATGITVSGAQSFKPVAWQLDNDGNKVNVDNRFATVT  LSATTGMKRGDKISFAGVKFLGQMAKNVLAQDATFSVVRVVDGTHVEITPKPVALDDVSLSPEQR  AYANVNTSLADAMAVNILNVKDARTNVFWADDAIRIVSQPIPANHELFAGMKTTSFSIPDVGLNG  IFATQGDISTLSGLCRIALWYGVNATRPEAIGVGLPGQTAGSGGGGSGAHIVMVDAYKPTKMEVQ  LVESGGGLVQAGGSLRLSCAASGRTFSSYAMGWFRQAPGKEREFVVAINWSSGSTYYADSVKGRF  TISRDNAKNTMYLQMNSLKPEDTAVYYCAAGYQINSGNYNFKDYEYDYWGQGTQVTVSSSGGGSG  GGDSATHIKFSKRDEDGKELAGATMELRDSSGKTISTWISDGQVKDFYLYPGKYTFVETAAPDGY  EVATAITFTVNEQGQVTVNGKATKG |

^a^ Underlined sequences were identified with high confidence.

**Table 3. Nucleic acid sequence of CP-ST_LS1 ^a^**

ATGGCACTGAACGAAGGCCAGATCGTCACCTTGGCAGTCGATGAGATAATTGAGACCATTAGTGCAATTACTCCTATGGCACAAAAGGCAAAGAAGTATACCCCACCGGCAGCAAGTATGCAGCGAAGTTCCAATACTATATGGATGCCTGTGGAGCAGGAGTCACCCACCCAGGAGGGATGGGATTTGACAGATAAG**GTTTGTTTCTGCTTCTACCTTTGATATATATATAATAATTATCATTAATTAGTAGTAATATAATATTTCAAATATTTTTTTCAAAATAAAAGAATGTAGTATATAGCAATTGCTTTTCTGTAGTTTATAAGTGTGTATATTTTAATTTATAACTTTTCTAATATATGACCAAAACATGGTGATGTTTAG**GCTACTGGGCTGCTTGAGTTGAATGTCGCAGTAAATATGGGTGAGCCCGACAATGACTTCTTTCAGTTGCGTGCAGACGACTTGAGGGATGAGACTGCATATCGACGACGAATCCAAAGCGCAGCCAGAAAACTCGCTAACAACGTCGAATTAAAGGTGGCCAACATGGCCGCAGAGATGGGGAGCCTCGTTATAACCTCTCCCGATGCCATAGGAACTAATACTGCAGACGCATGGAATTTCGTTGCAGACGCCGAGGAGATAATGTTCAGCAGGGAGCTGAACAGGGATATGGGTACTTCTTATTTTTTCAATCCTCAAGATTATAAAAAAGCAGGCTATGATCTCACAAAAAGGGATATATTTGGCCGTATACCCGAAGAAGCTTACAGGGACGGAACCATTCAGCGACAAGTGGCAGGATTCGATGATGTTTTACGATCCCCCAAGCTTCCTGTCCTGACTAAATCCACGGCAACGGGGATAACGGTCTCAGGGGCCCAGTCATTCAAACCAGTGGCCTGGCAACTTGATAACGATGGGAACAAAGTTAACGTTGATAACCGTTTTGCCACTGTGACCCTCAGTGCCACCACTGGTATGAAAAGGGGGGATAAAATAAGCTTTGCTGGCGTCAAGTTTCTTGGACAAATGGCCAAGAATGTGTTGGCCCAGGACGCCACCTTTTCAGTGGTGCGAGTGGTCGACGGAACGCACGTAGAAATCACCCCCAAACCAGTCGCTTTAGATGATGTATCCCTTTCCCCAGAGCAAAGGGCATACGCTAACGTAAACACATCCCTTGCAGATGCCATGGCTGTAAACATACTCAACGTGAAGGATGCCAGAACCAACGTCTTCTGGGCAGATGATGCCATCAGGATTGTTTCCCAGCCGATTCCCGCAAACCATGAACTTTTCGCCGGAATGAAAACGACTTCTTTTTCTATACCGGACGTTGGCCTGAACGGTATTTTTGCCACCCAGGGCGATATTAGCACTTTAAGTGGTCTGTGTCGAATCGCCTTGTGGTACGGAGTCAATGCTACTAGACCGGAGGCCATAGGGGTTGGCTTGCCAGGTCAAACTGCCTAATAG

^a^ **Bold and underlined** sequence represents the ST_LS1 intron. Red sequence represents stop codons, with the stop codon contained within ST_LS1 also shown.

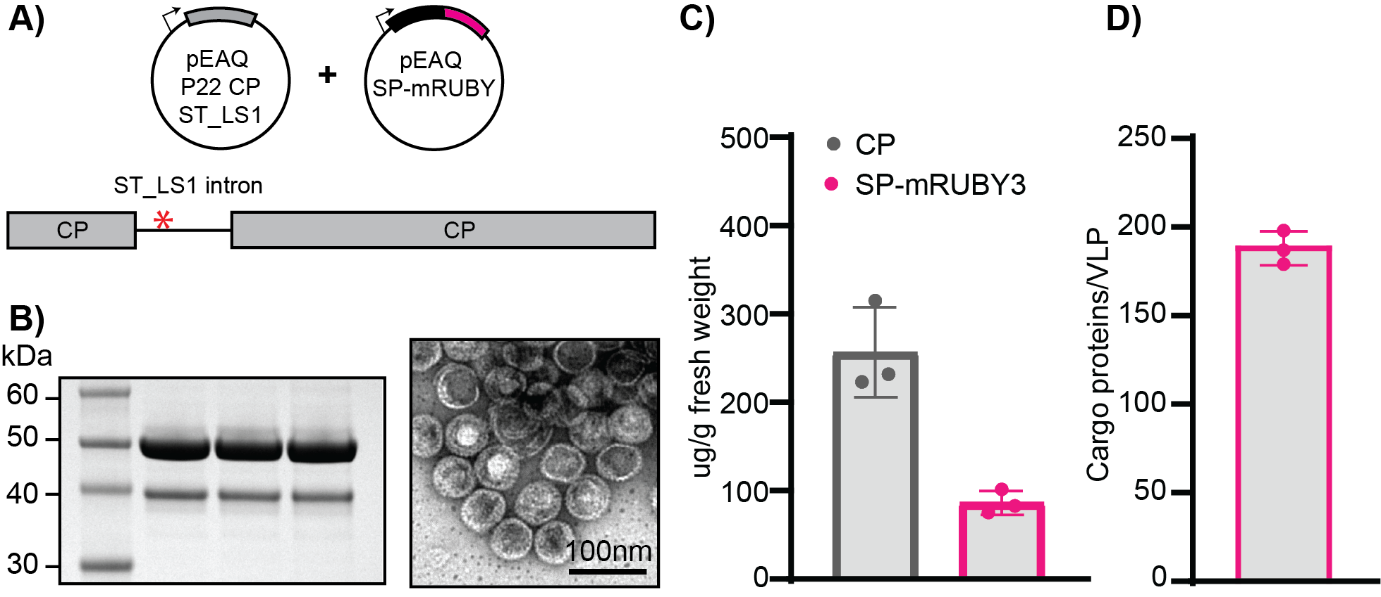

**Figure S1. Intron-containing CP expression.**

**A)** Cartoon illustration of the two pEAQ vectors containing CP-ST_LS1 and SP-mRUBY3 respectively. The ST_LS1 intron is inserted such that, if not spliced, protein translation will be terminated with a stop codon after 73 amino acids. Should the intron be spliced, normal CP translation will be achieved. **B)** Coomassie stained gel (SDS-PAGE) of ultracentrifuged protein extracts n=3 transient expressions. The presence of a both bands corresponding to the full-length CP and SP-mRUBY3 suggests correct splicing of the ST_LS1 intron. A representative negative stain TEM from one of the replicates is also shown, with the 100 nm scale bar displayed. **C)** Average calculated yield of purified CP and SP-mRUBY3 in ug/g plant fresh weight (n=3). **D)** Average calculated SP-mRUBY3 cargo loading per VLP (assuming 420 CP).

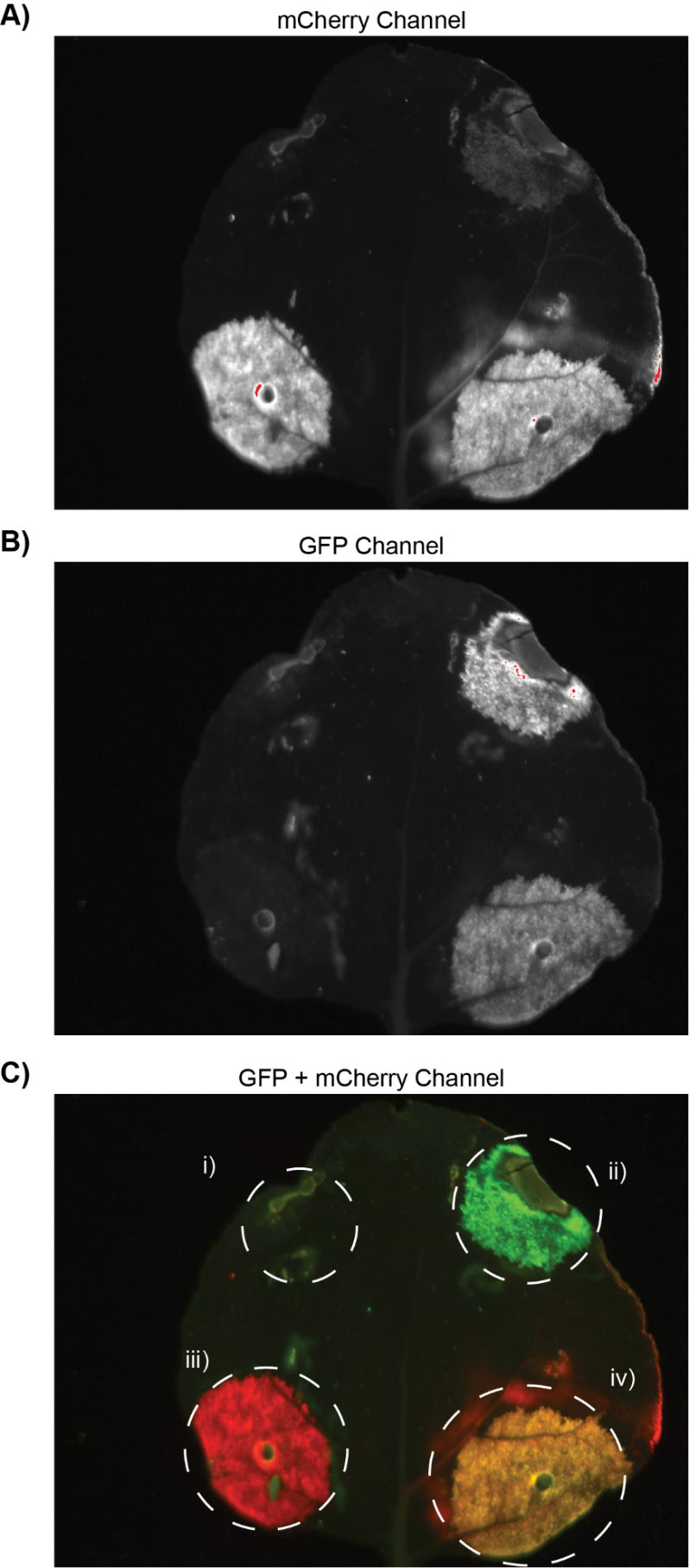

**Figure S2. Fluorescent imaging of infiltrated leaf tissue.** Images of a single leaf infiltrated with (i) CP alone, (ii) CP and SP-mCLOVER3, (iii) CP and SP-mRUBY3, and (iv) CP, SP-mCLOVER3, and SP-mRUBY3. Images corresponding to the red fluorescence channel, green fluorescence channel, and a merge of both channels are shown in **A)**, **B)**, and **C)** respectively.

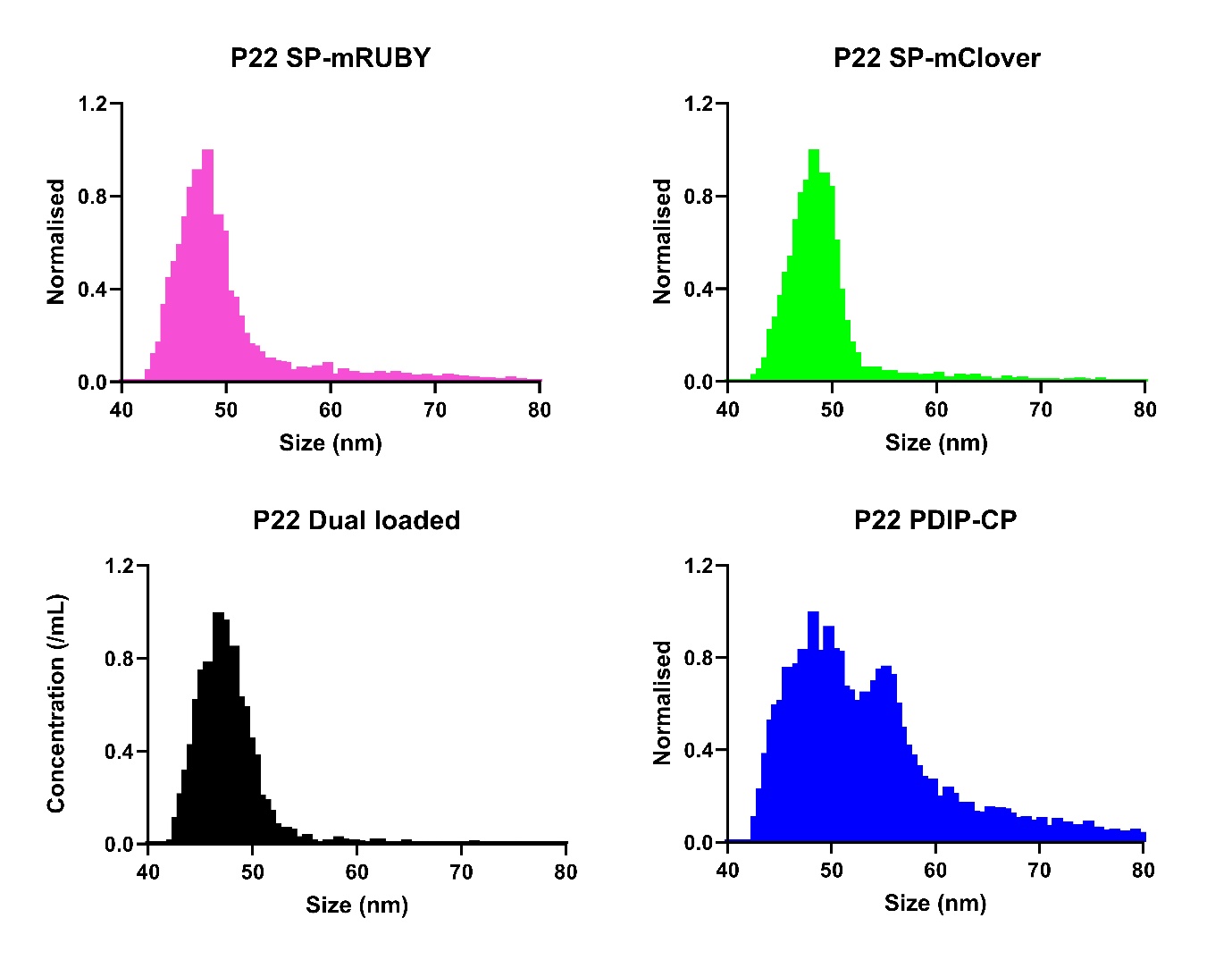

**Figure S3. Nano flow cytometry of purified VLPs.** Nano flow cytometry spectra for purified VLPs packaged with SP-mRUBY3 (pink), SP-mCLOVER3 (green), co-loaded with both SP-mRUBY3 and SP-mCLOVER3 (black), and the PDIP-CP fusion (blue).

**z**
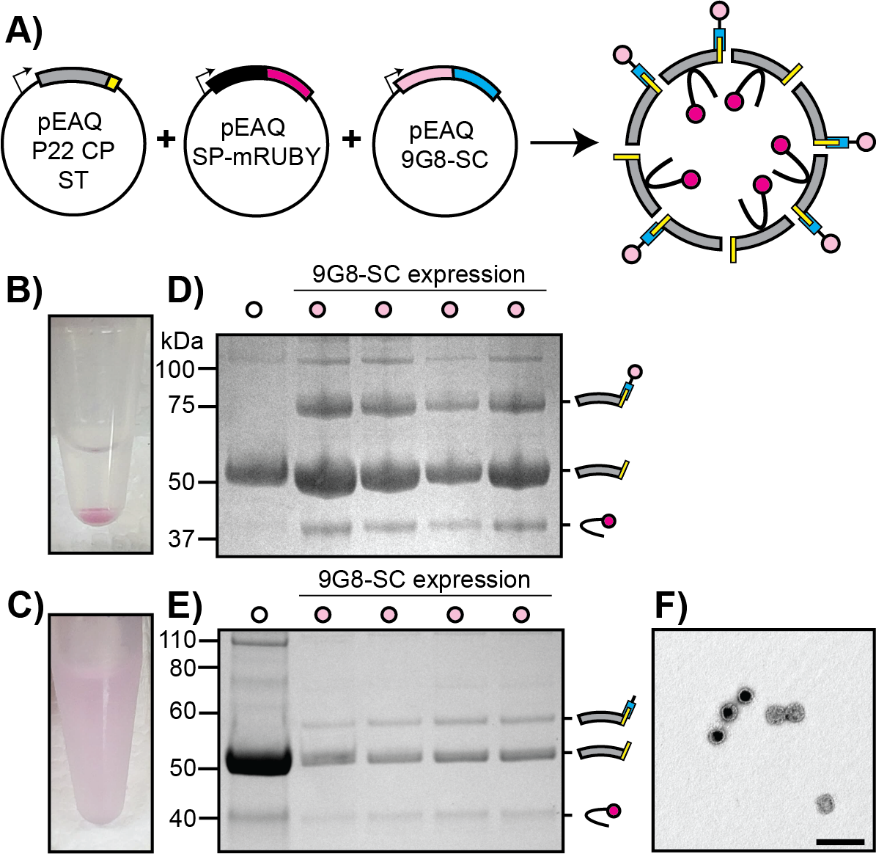

**Figure S4. Exploring display of 9G8 nanobody. A)** Cartoon illustration of pEAQ vectors with 9G8-SC nanobody co-expression. **B)** Insoluble pink pellet, collected from top of PD10 desalting column after failing to pass through filter. Resuspended in PBS. **C)** Soluble pink protein desalted through PD10 column. **D, E)** Coomassie stained SDS-PAGE of purified VLP-ST (left) and VLPs co-expressed with 9G8-SC (n=4). (**D)** is from of the insoluble pink protein pellet and **(E)** is from soluble pink protein. A clear difference in VLP products is seen, with the insoluble sample **(D)** containing full-length 9G8-SC fused to P22 CP-ST and the soluble sample (**E)** only containing the CP-ST/SC cleavage product. **F)** Negative stain TEM of the soluble sample (scale bar indicates 200nm). The insoluble sample could not be visualised by TEM.

Following purification by ultracentrifugation and desalting on PD10 column, it was identified that although the column elution was pink, there remained pink aggregates which had not passed through the column. Removal of these aggregates from the top of the column and dilution in PBS clearly showed insoluble pink protein compared to that collected from the elution (Figure S4B and S4C respectively). Analysis of both of these proteins by denaturing SDS-PAGE produced unexpected results. The insoluble pellet showed very clean protein species corresponding to P22 VLPs with CP-ST functionalised with 9G8-SC (Figure S4D). The protein species ~75 kDa was confirmed to be the CP-ST/SC-9G8 conjugate by tryptic digest LC-MS/MS (Table S2). No breakage product could be seen on this gel. Comparatively, the soluble desalted sample did display the CP-ST/SC breakage product, but not full-length fusion containing 9G8 (Figure S4E). The soluble sample could be imaged by negative stain TEM (Figure S4F) to show correctly formed VLPs, however the insoluble sample could not be visualised.

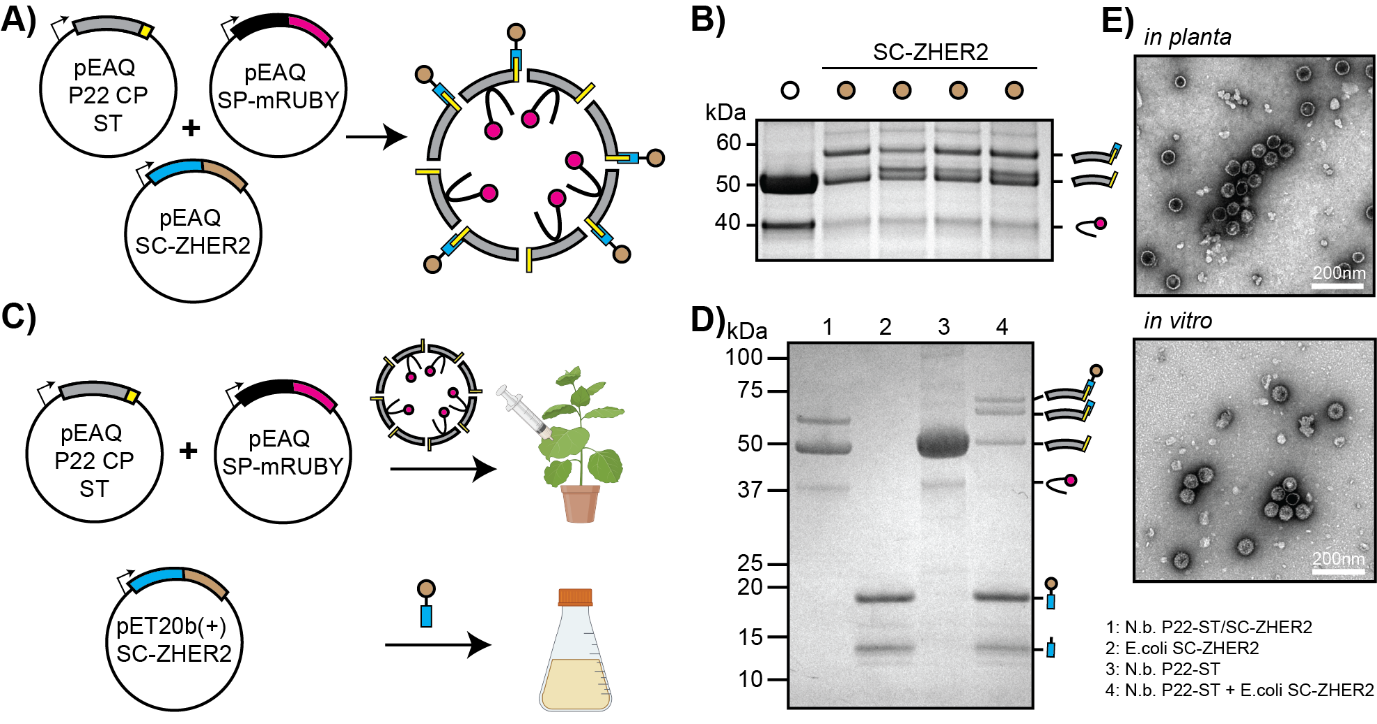

**Figure S5. Exploring display of ZHER2 affibody. A)** Cartoon illustration of pEAQ vectors with SC-ZHER2 affibody co-expression. **B)** Coomassie stained SDS-PAGE of purified VLP-ST (left lane) and VLPs co-expressed with SC-ZHER2 (n=4). Negative stain TEM of VLPs co-expressed with SC-ZHER2. **C)** Illustration of vector expression in *N. benthamiana* and in *E. coli* to assess SC-ZHER2 stability. **D)** Coomassie stained SDS-PAGE with samples from (1) SEC purified CP-ST/SC-ZHER2 expression from *N. benthamiana*, (2) HisTrap purified SC-ZHER2 from *E. coli*, (3) ultracentrifuged P22 VLP-ST from *N. benthamiana*, and (4) overnight *in vitro* incubation of proteins from lane (2) and lane (3). **E)** Negative stain TEM of VLPs purified from *N. benthamiana* following co-expression of CP-ST, SP-mRUBY3, and SC-ZHER2 (top) and of VLPs from *in vitro* mixing experiment (bottom) from lane (4) of figure (D).

Co-expression of SP-mRUBY3, CP-ST, and SC-ZHER2 in *N. benthamiana* (Figure S5A) failed to generate VLPs that displayed full-length SC-ZHER2. SDS-PAGE and tryptic digest of the ~58 kDa band confirmed the formation of an isopeptide bond between CP-ST and SC, and TEM confirmed correct assembly of VLPs (Figure S5B). However, the full-length SC-ZHER2 fusion (~68.6 kDa expected molecular weight) was not detected, and subsequent size exclusion chromatography (SEC) of all four replicate expressions pooled together showed the presence of only three protein species on SDS-PAGE (Figure S5D, lane 1) that corresponded to SP-mRUBY3, CP-ST and CP-ST/SC, in increasing molecular weight respectively. These results suggest that the ZHER2 affibody, which has not previously been produced in plants, might be liable to degradation in *N. benthamiana*. To investigate this potential limitation, the SC-ZHER2 fusion was cloned into a modified pET20b(+) *E. coli* expression vector (without PelB periplasmic signal peptide) for production in the cytosol of BL21(DE3) cells (Figure S5C), with the aim of replicating the successful *in vitro* conjugation of SC-ZHER2 to SpyTagged-P22 VLPs that was reported (Kim et al., 2019).

HisTrap purification of SC-ZHER2 from *E. coli* produced two clean protein species (Figure S5D, lane 2), corresponding to the expected molecular weight for both the full-length SC-ZHER2 and SC alone. The SC-ZHER2 purified from *E. coli* was incubated with SpyTagged-P22 VLPs purified from *N. benthamiana* (Figure S5D, lane 3), overnight at 4ºC. SDS-PAGE analysis of the *in vitro* conjugation reaction revealed the presence of an additional higher-molecular weight protein species that corresponded to the full-length SC-ZHER2 (Figure S5D, lane 4). Correct assembly of the P22 VLPs was confirmed by negative stain TEM (Figure S5D).
